# Spectral organ fingerprints for intraoperative tissue classification with hyperspectral imaging

**DOI:** 10.1101/2021.11.24.469943

**Authors:** A. Studier-Fischer, S. Seidlitz, J. Sellner, M. Wiesenfarth, L. Ayala, B. Özdemir, J. Odenthal, S. Knödler, K.F. Kowalewski, C.M. Haney, I. Camplisson, M. Dietrich, K. Schmidt, G.A. Salg, H.G. Kenngott, T.J. Adler, N. Schreck, A. Kopp-Schneider, K. Maier-Hein, L. Maier-Hein, B.P. Müller-Stich, F. Nickel

**Affiliations:** Department of General, Visceral, and Transplantation Surgery, Heidelberg University Hospital, Heidelberg, Germany; Division of Computer Assisted Medical Interventions, German Cancer Research Center (DKFZ), Heidelberg, Germany; HIDSS4Health – Helmholtz Information and Data Science School for Health, Karlsruhe, Heidelberg, Germany; Division of Medical Image Computing, German Cancer Research Center (DKFZ), Heidelberg, Germany; Division of Biostatistics, German Cancer Research Center (DKFZ), Heidelberg, Germany; Department of Urology, Medical Faculty of Mannheim at the University of Heidelberg, Mannheim, Germany; Division of Biology and Biological Engineering, California Institute of Technology, Pasadena, USA; Department of Anesthesiology, Heidelberg University Hospital, Heidelberg, Germany; Department of Anesthesiology and Intensive Care Medicine, Essen University Hospital, Essen, Germany; Faculty of Mathematics and Computer Science, Heidelberg University, Heidelberg, Germany

**Keywords:** hyperspectral imaging, multispectral imaging, organ classification, tissue classification, organ identification, tissue identification, translational research, porcine model, machine learning, deep learning, linear mixed model, surgery, surgical data science

## Abstract

Visual discrimination of tissue during surgery can be challenging since different tissues appear similar to the human eye. Hyperspectral imaging (HSI) removes this limitation by associating each pixel with high-dimensional spectral information. While previous work has shown its general potential to discriminate tissue, clinical translation has been limited due to the method’s current lack of robustness and generalizability. Specifically, it had been unknown whether variability in spectral reflectance is primarily explained by tissue type rather than the recorded individual or specific acquisition conditions. The contribution of this work is threefold: (1) Based on an annotated medical HSI data set (9,059 images from 46 pigs), we present a tissue atlas featuring spectral fingerprints of 20 different porcine organs and tissue types. (2) Using the principle of mixed model analysis, we show that the greatest source of variability related to HSI images is the organ under observation. (3) We show that HSI-based fully-automatic tissue differentiation of 20 organ classes with deep neural networks is possible with high accuracy (> 95 %). We conclude from our study that automatic tissue discrimination based on HSI data is feasible and could thus aid in intraoperative decision making and pave the way for context-aware computer-assisted surgery systems and autonomous robotics.

## 1 Background

Discrimination of tissue conditions, pathologies and critical structures from healthy surrounding tissue during surgery can be challenging given the fact that different body tissues appear similar to the human eye. While conventional intraoperative imaging is limited by mimicking the human eye, hyperspectral imaging (HSI) removes this arbitrary restriction of recording only red, green and blue (RGB) colors. HSI works by assigning each pixel of a conventional two-dimensional digital image a third dimension of spectral information. The spectral information contains the wavelength-specific reflectance intensity of every pixel. This results in a three-dimensional datacube with two spatial dimensions (*x, y*) and a third spectral dimension (*λ*). HSI has found application in diverse fields such as geology and maritime studies, agriculture, food industry, automated waste sorting [1, 2] and has recently been used during a NASA space mission on Mars.

Over the last few years, there have been extensive efforts to implement HSI technology in healthcare. Examples of potential future clinical applications comprise the objective evaluation of tissue oxygenation and blood perfusion [3-6], inflammation and sepsis [7] or malignancy [8] as well as computer-assisted decision-making and automated organ identification [9]. These have the potential to support future developments such as intraoperative cognitive assistance systems or even automatization of robotic surgery. Despite the promising research, clinical translation of HSI-based automatic tissue differentiation has not yet been achieved. This may be attributed to a current lack in robustness and generalizability, which are the most important requirements for clinical application. In this regard, several open research questions remain. Specifically, variability of HSI measurements may result from the inherent differences between multiple tissue types under observation (desired effect), but also from inter-subject variability or variability in image acquisition conditions (both undesired). We are not aware of any prior work that has systematically investigated this important topic and we ultimately aim to provide a thorough understanding of hyperspectral organ data, illustrate the potential of HSI-based analyses and present solid baseline data that further studies can build upon.

## 2 Results

For automatic tissue characterization based on HSI data, the following two properties are highly desirable: First, spectra corresponding to different organs should differ substantially from each other. And second, spectra of the same organ should be relatively constant across image acquisition conditions and individuals. With this in mind and given the gap in literature pointed out in the section above, the contribution of this work is threefold:

- Spectral fingerprints: We present the first comprehensive analysis of spectral tissue properties for a wide range of physiological organs and tissue types. Based on 9,059 images of 46 pigs and 17,777 annotations, we generate specific spectral fingerprints for a total of 20 organs.
- Variance analysis: We show that the greatest part of spectral variance can be explained by organ differences.
- Machine learning-based organ and tissue classification with HSI: We demonstrate that a neural network can distinguish between organ classes with high accuracy (> 95%), suggesting that HSI has high potential for intraoperative organ and tissue discrimination

### 2.1 Different organs feature unique spectral fingerprints

This project provides insight on the spectral reflectance of 20 porcine organs in a total number of 9,059 images within 46 animals (**Figure 1**). Our data shows that different organs feature characteristic spectra, which motivated us to refer to them as organ “fingerprints”. As seen in the gray pig-specific reflectance curves in **Figure 1**, variation in the spectral measurements may result not only from the organ, but also from the individuals and/or the specific measurement conditions. A key aim of this work was therefore to quantify the effect of the different sources of variation.

**Figure 1.**
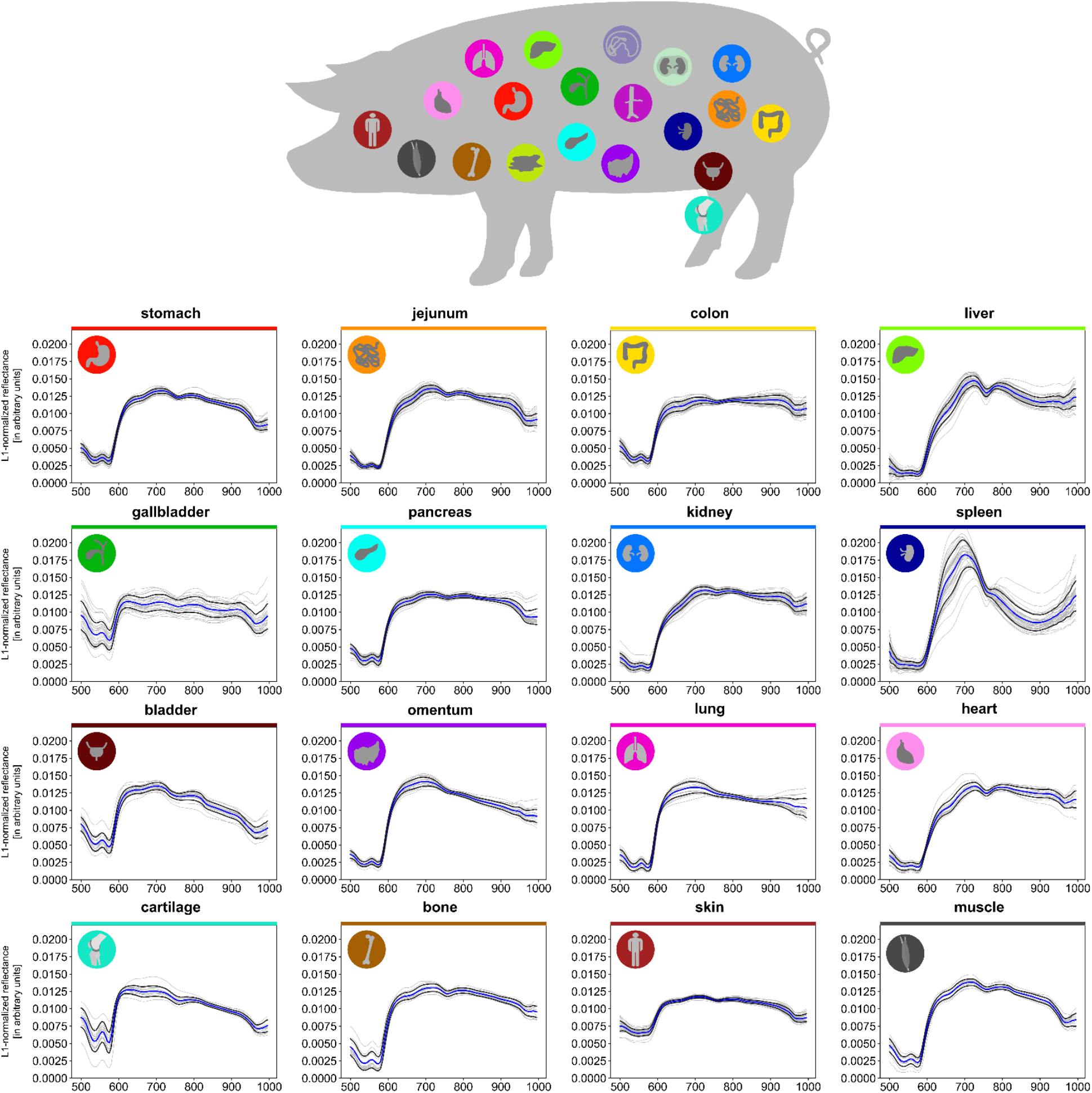
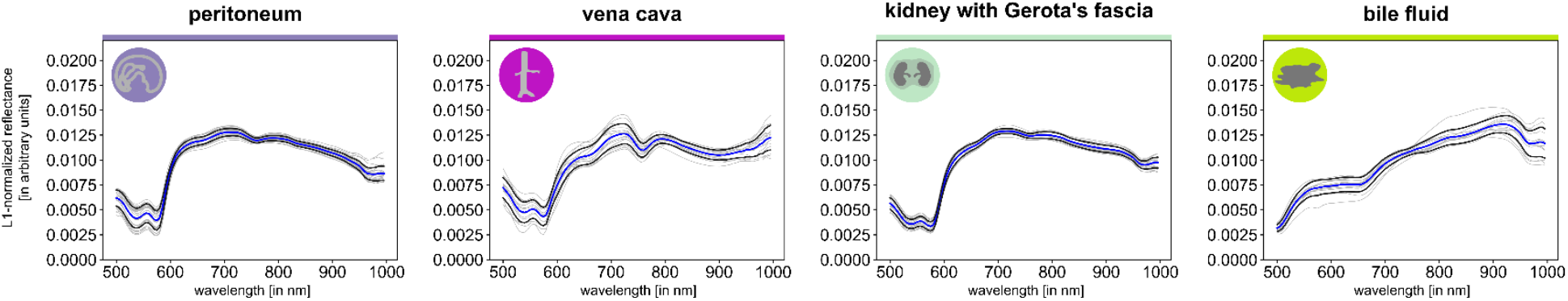
Tissue atlas comprising spectral fingerprints of 20 abdominal and thoracic organs. Stomach (A=39; n=849), jejunum (A=44; n=1,546), colon (A=39; n=1,330), liver (A=41; n=1,454), gallbladder (A=28; n=526), pancreas (A=31; n=530), kidney (A=42; n=568), spleen (A=41; n=1,353), bladder (A=32; n=779), omentum (A=23; n=570), lung (A=19; n=652), heart (A=19; n=629), cartilage (A=15; n=586), bone (A=14; n=537), skin (A=43; n=2,158), muscle (A=15; n=560), peritoneum (A=28; n=2,042), vena cava (A=15; n=353), kidney with Gerota’s fascia (A=18; n=393), bile fluid (A=13; n=362). A indicates the number of animals; n indicates the number of measurements in total. Graphs depict mean reflectance (L1-normalized on pixel-level) of individual pigs (gray) as well as overall mean (blue) ±1 standard deviation (SD) (black) with wavelengths from 500 to 995 nm on the x-axis and reflectance in arbitrary units on the y-axis.

### 2.2 Spectral similarity between organs is heterogeneous

In order to illustrate the HSI variability resulting from individuals and measurement conditions, we applied t-distributed Stochastic Neighbor Embedding (t-SNE) [10] to our data (**Figure 2**). It shows that while certain tissue types such as spleen and liver form highly isolated clusters, other organs such as stomach, pancreas and jejunum have a tendency to overlap, indicating lower distinguishability.

**Figure 2.**
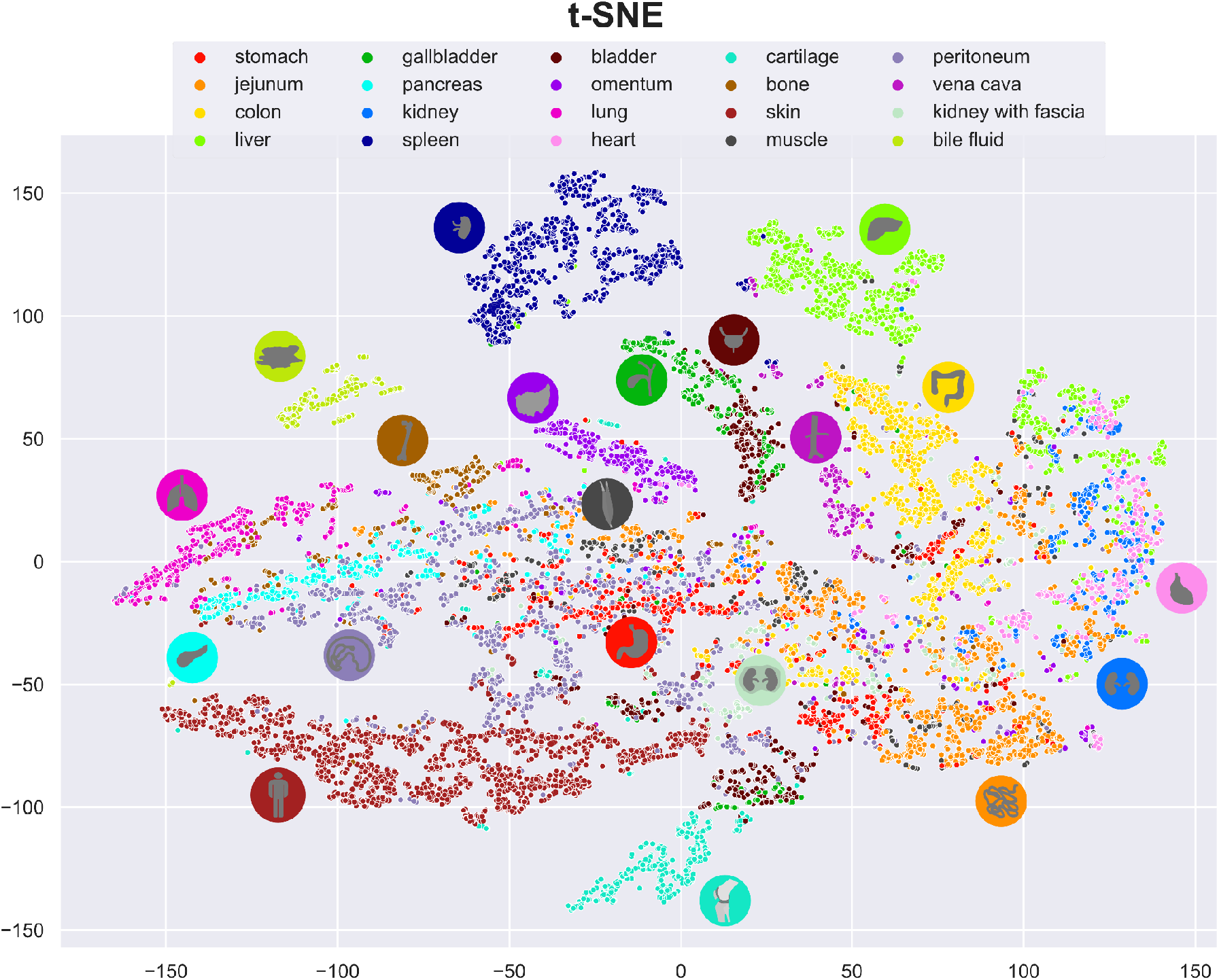
Visualization of spectral similarity with t-SNE as a non-linear multi-dimensionality reduction tool; one point represents the median spectrum within a region of interest (ROI) of one organ in one image of one pig. It can be seen that organs such as spleen and liver form isolated clusters while other organs such as jejunum greatly overlap with the rest.

### 2.3 Organ is the most influential factor on the reflectance spectrum

To quantify the effect of different sources of variation, we applied linear mixed models on a highly standardized subset of data obtained from 11 pigs (within P36 to P46 as illustrated in **Supplementary Figure 1**). The analysis was performed at first for all organs (**Figure 3**) and subsequently stratified by organ (**Figure 4**). In the analysis for all organs, at each wavelength the proportion of explained variation [11] in observed reflectance was decomposed into the components “organ”, “pig”, “angle”, “image” and “repetition”, where “angle” describes the proportion of variation explained by the angle between the organ surface and the camera optical axis, “image” describes the proportion of variation explained by different measurements taken from different organ positions in the same individual or variations in the annotated areas, and “repetition” describes the proportion of explained variation by multiple recordings of the same image under identical measurement conditions.

**Figure 3.**
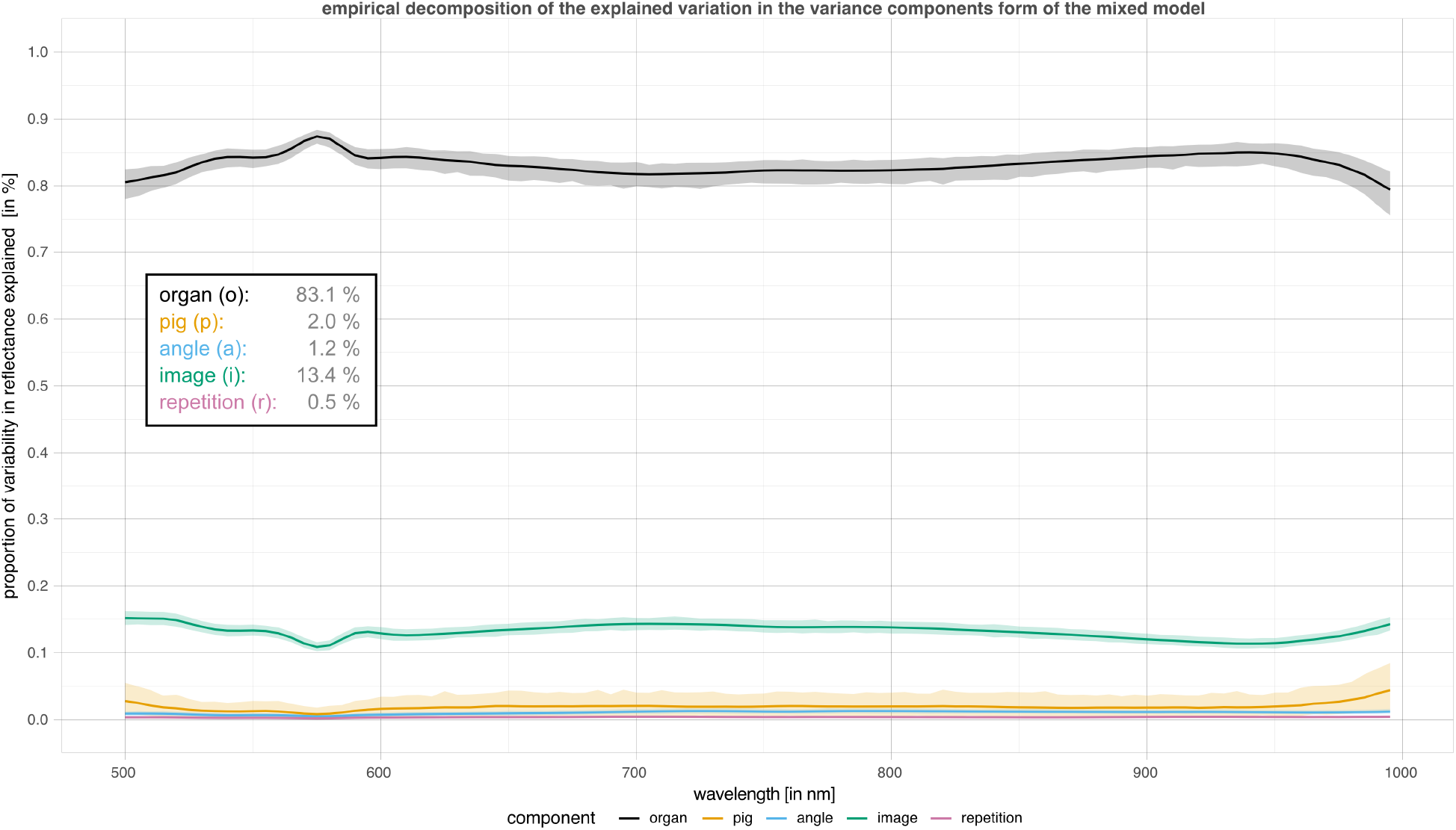
Sources of variation of hyperspectral data. (Proportion of) variability in reflectance explained by each factor using linear mixed models. Factors include “organ”, “pig”, “angle”, “image” and “repetition”. For each recorded wavelength, an independent linear mixed model was fitted with fixed effects for the factors “organ” and “angle” as well as random effects for “pig” and “image”. Variation across repetitions was given by the residual variation. The greater the proportion of variability for “organ”, the more reflectance can be seen as organ-characteristic. Shaded areas depict 95 % (pointwise) confidence intervals based on parametric bootstrapping. The numbers represent the median across wavelengths.

**Figure 4.**
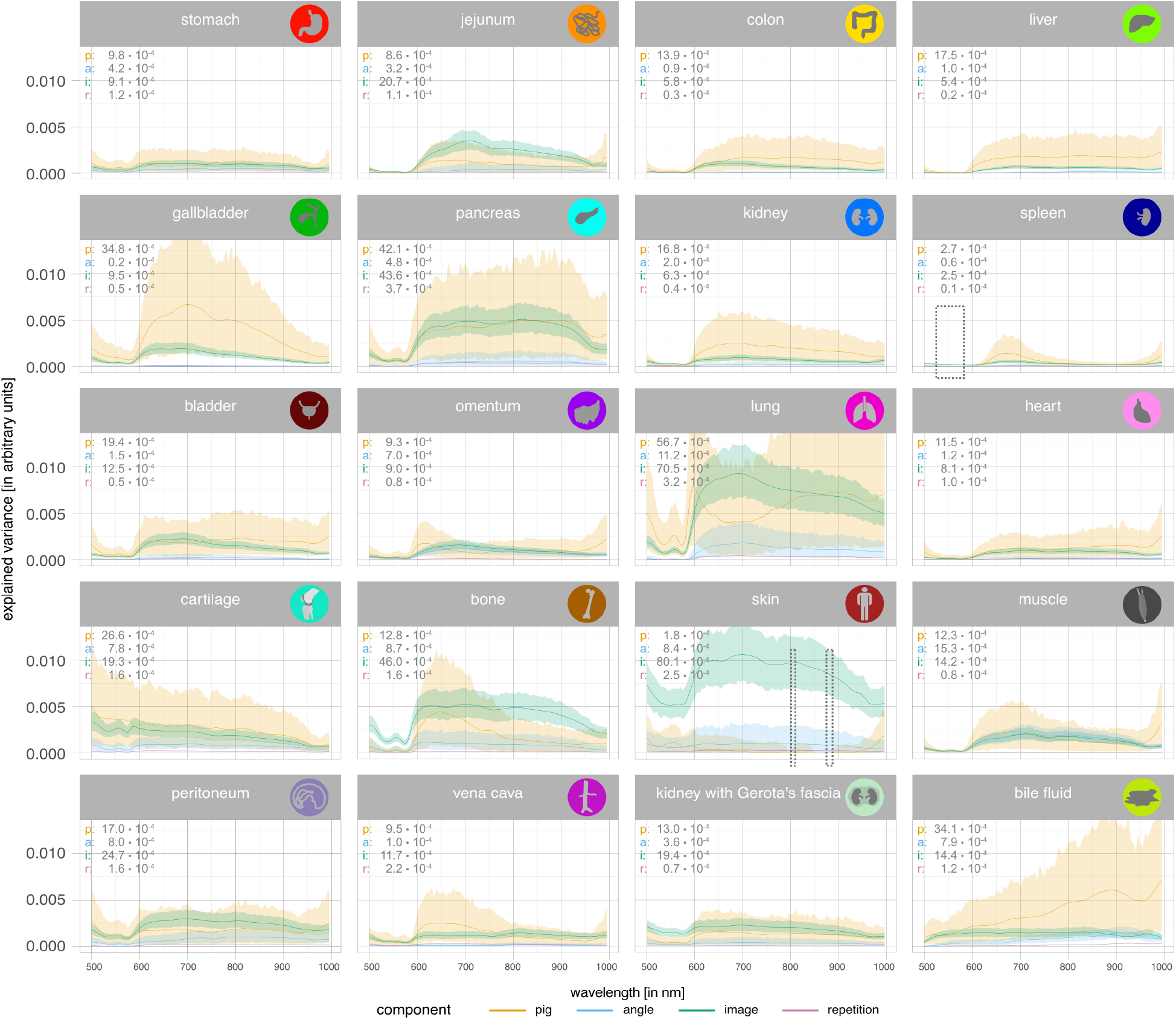
Sources of variation of hyperspectral data stratified by organ. Explained variation analysis stratified by organ using linear mixed models. For each organ and wavelength, independent linear mixed models were fitted with fixed effects for “angle” and random effects for “pig” and “image”. Variation across repetitions is given by the residual variation. Shaded areas depict 95 % (pointwise) confidence intervals based on parametric bootstrapping. The numbers on each subplot represent the median across wavelengths. Model fit was singular for skin and spleen for some wavelengths (spleen: 520–585 nm; skin: 800–810, 845 and 870–885 nm) for which curves were linearly interpolated (dashed boxes).

Our analysis suggests that the main influencing factor on HSI data variation across wavelengths was the factor “organ” with an average proportion of explained variability of 83.3 %. The factor “image” explained 13.2 % of the variation on average while the other factors only explained negligible variation with 2.0 % for “pig”, 1.1 % for “angle” and 0.4 % for “repetition”. This suggests that HSI data is characteristic of organs much more than of the subjects under observation or other influencing factors. The percentage to which variance in reflectance was explained by the components was not constant, but varied slightly through different parts of the recorded electromagnetic spectrum.

When stratifying by organ, the variance in reflectance was decomposed into the same components except “organ”. According to **Figure 4**, “angle” and “repetition” explain a negligible portion of the variance in all organs. For “pig” and “image”, differences between organs are present. For organs where all lines are close to zero (e.g. spleen), there is essentially no heterogeneity in reflectance between different images and pigs, thus these organs show the most pronounced organ-characteristic spectral signatures. On the other hand, organs with greater levels of explained variance for the components “pig” and “image” consequently had less organ-characteristic spectral signatures, such as lung with a comparatively high mean value of 0.0054 for variation explained by “pig”. Organ classes with the highest cumulative levels of variance curves explained by factors other than “organ” and therefore the least organ-characteristic spectral signatures across observations were lung and skin (**Supplementary Text 4** and **Supplementary Table 1**).

For some organs, such as the gallbladder, reflectance varied strongly between pigs (value for “pig” comparatively high), but little within a pig (value for “image” comparatively low). Thus, reflectances measured for gallbladders were heterogeneous across individual pigs. On the other hand, for other organs, heterogeneity within pigs (i.e. between images of the same pig) was much larger (value for “image” high) than between pigs (value for “pig” low), e.g. for skin. Thus, reflectance measured for skin tends to be homogeneous across individual pigs, but a single image of skin may be unreliable due to the heterogeneity within one pig.

### 2.4 Machine learning can leverage spectral information to classify tissue with high accuracy

A deep learning-based approach was used to classify the annotations of 20 organ classes from the spectra presented above with an average accuracy of 95.4 % ± 3.6 % across pigs on a hold-out test set. Misclassification only occurred for 486 out of 9,895 annotations in the test set (**Figure 4**). While 16 out of the 20 organ classes were classified with an average sensitivity of ≥ 90 % across all pigs, the smallest average sensitivity across pigs was obtained for the organ classes gallbladder (74.0 %) and heart (73.9 %), which were on average across pigs most often confused with bladder and kidney, respectively.

**Figure 4.**
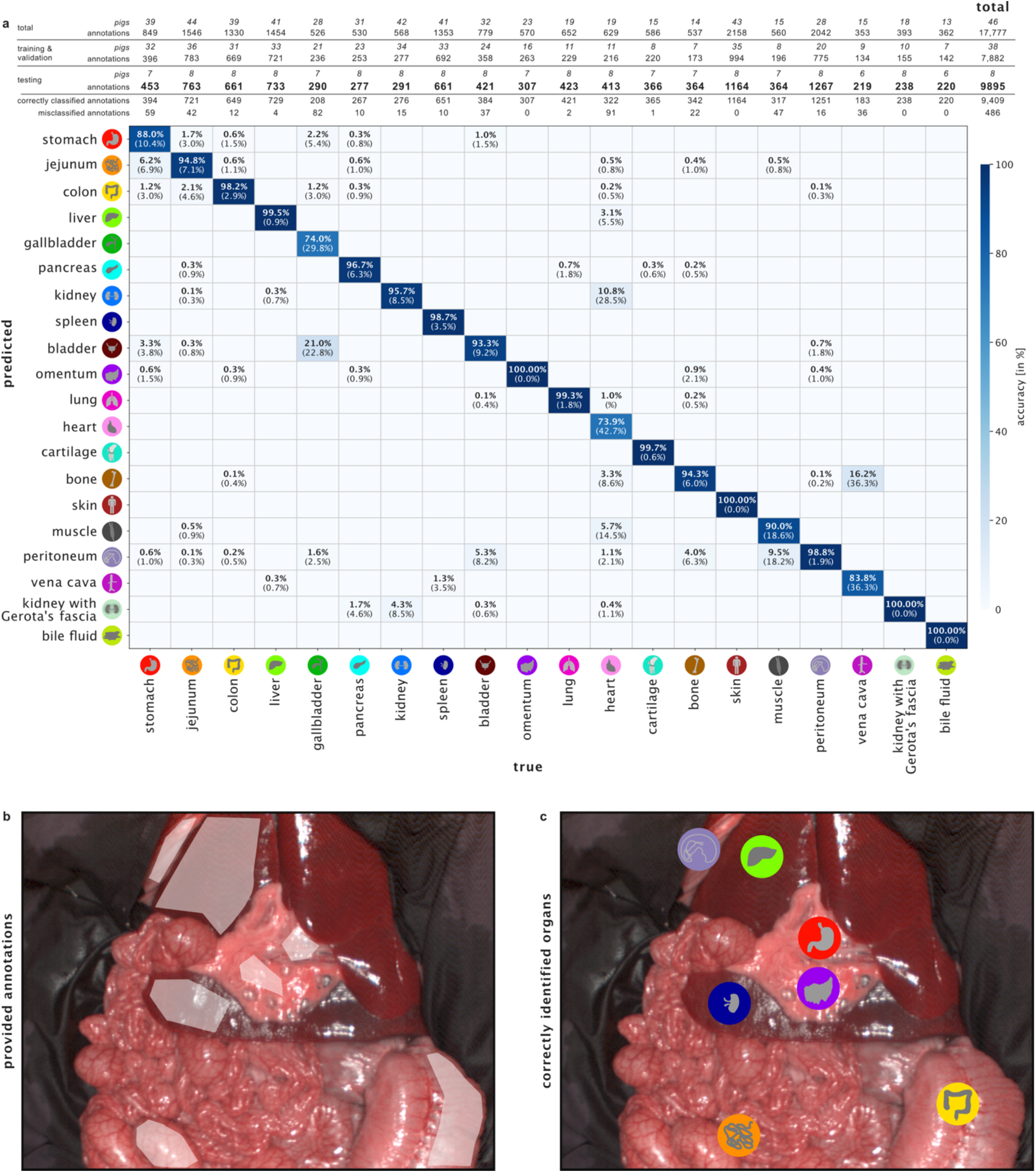
Results of deep learning-based organ classification. **a**, confusion matrix which was generated for a hold-out test set comprising 9,895 annotations from 5293 images of 8 pigs that were not part of the training data. Confusion matrices were calculated and column-wise normalized (i.e. divided by the column sum) per pig based on the absolute number of (mis-)classified annotations. These normalized confusion matrices were averaged across pigs while ignoring non-existent entries (e.g. due to missing organs for one pig). Each value in the matrix thus depicts the average fraction of annotations which were labeled as the column class and predicted as the row class. Numbers in brackets depict the standard deviation across pigs. Zero values are not shown in the confusion matrix in order to improve visibility. Since multiple organs can appear on the same image, the number of annotations exceeds the number of images. **b**, exemplary image with multiple organ annotations by an expert. **c**, organs classified through deep learning.

## 3 Discussion

Visual discrimination and evaluation of biological tissue is not trivial, as different tissues and body structures often appear similar to the human eye. Because conventional optical imaging during surgery only differentiates red, green and blue by mimicking human vision, its intraoperative benefit is sometimes limited. HSI, not being subject to this restriction and encompassing significantly more information, is an exceptional imaging modality with great potential for tissue identification and evaluation. Although its current use in medicine is on a constant rise, the full potential of this imaging modality has not been exploited. This may be attributed to open research questions concerning robustness and generalizability of HSI data.

Structural properties of tissue cause differences in spectral characteristics that might be significant enough for use in proper organ differentiation and other clinical applications. However, existing literature on spectral measurements has mainly focused on specific biological pigments such as hemoglobin, porphyrin and melanin [12, 13], and has hardly addressed the complexity of spectral characteristics across various tissues and organs. Moreover, past works have often focused on optical scattering instead of reflectance or absorbance and therefore provided data of limited practical applicability, have investigated ex-vivo material, featured low wavelength resolution or performed measurements with incompatible, incomparable and outdated technology that does not allow for comparison between different studies [14]. Despite its necessity as a foundation for advanced studies, as of yet, no systematic database or investigation of reflectance spectra for a variety of physiological organs in a larger cohort existed.

Categorical requirements for such a spectral medical HSI database serving as a reference work are precision, uniformity and comparability of the measuring device, which has been demanded by the HSI community in previous years [15]. While in former decades HSI could not be found in medicine, there have been extensive efforts to implement this technology in healthcare over the last years. However, most of the initially developed HSI systems were self-made prototypes and homebuilt solutions from various institutions all over the world, varying in spectral resolution and range as well as utilized detectors and optical components [15-25]. While highly interesting insights for various medical applications were obtained with these provisional solutions, they were lacking standardization and reproducibility [15], rendering sustainable large clinical trials and systematic multicenter research impossible. A great variety of devices could be observed in terms of spectral resolution, detectors, dispersive devices and spectral regions covered by different devices reaching from 200 nm up to 2500 nm [15].

The HSI camera system used in this project is the first commercially available and medically certified system meeting the aforementioned demands. While previous and less standardized HSI systems were efficient for the investigation of specific and isolated research questions, reproducibility and generalizability of commercially available systems noticeably promoted an increase in research efforts regarding HSI. An indicator of these increased research efforts can be seen in the rise of the number of research projects over the last few years including animal studies with rats [26] and pigs [6, 27-29], conference papers [30, 31], narrative reviews [32-34] and other publications [35, 36]. With this new system and its advantages, special focus has again been put on early clinical trials with explorative character [1, 37-52]. However, there are novel possibilities that have not been exploited yet. These primarily include spectral characterization of biological tissue and the complementation of a large medical HSI database with machine learning and deep learning. Some studies have already spectrally characterized single aspects of biological tissue such as the differences between specific cancer entities and their related physiological tissue [53]. However, these studies were most often conducted with non-reproducible setups or sometimes done in-vitro without measuring tissue perfusion, which might be acceptable for specific bradytrophic tissues, but leads to limitations in applicability to the majority of typically well-perfused organs [54]. Moreover, most of the existing studies so far only highlight specific medical aspects and do not sufficiently broaden the general understanding of spectral tissue characteristics.

The principles of spectral tissue differentiation have already successfully been proven, however only in laparoscopic surgery with sparse multispectral information and, most importantly, in fewer organ classes [55, 56]. The question driving the present study was whether these spectral differences would be strong enough to be detected by an HSI system and subsequently consistent enough to characterize organs and make organ differentiation feasible.

For the very first time, HSI was applied with the aim of (1) systematically characterizing spectral properties of different tissue types in a porcine model, (2) analyzing to which extent these spectra are influenced by organ or tissue type compared to undesired effects such as inter-subject variability and variations in image acquisition conditions and (3) demonstrating that automatic machine learning-based tissue classification even with an unusually high number of classes can be achieved with high accuracy. A total number of 20 different porcine organs were recorded with HSI. The resulting database comprises 9,059 recordings with 17,777 annotated organ regions.

Spectral fingerprints of these organs were extracted in **Figure 1** and t-SNE was chosen to visually assess the distinguishability of the respective HSI spectra (**Figure 2**). While Euclidean distances have to be interpreted cautiously in illustrations from high-dimension reduction tools, clustering and overlap give a good hint at the differentiability of the underlying spectra. It was now essential to evaluate to what extent differences in reflectance could be attributed to the organ or alternatively to the individual pig or noise from other defined and undefined factors as this would determine the general utility of HSI data.

Linear mixed models could show that the largest proportion of the spectral reflectance variability was attributed to the factor “organ” instead of “pig”, “angle”, “image” and “repetition”. This suggests that contributions from inter-individual differences and image acquisition conditions were dominated by organ differences. While image acquisition conditions such as illumination were highly standardized, artificial over-standardisation was consciously avoided in order to still comply with conditions in the real operative room. Of the other factors accounting for spectral reflectance variability, “image” was the most relevant one, which indicates, that different regions on the same organ have spectral differences. Possible explanations for this finding are inhomogeneous distribution of connective tissues, blood vessels and fibrosis within each organ, different levels of contained blood volumes due to tension on the tissue surface or peristalsis. This insight explicitly for the influencing factor “image” is highly relevant when considering possible real-life intraoperative applications and trials, as it - depending on the depth of the analysis - implies the necessity to record different areas of the organ under investigation as we did.

A machine learning algorithm had an average accuracy of over 95 % in an independent test set for identifying organ classes on pre-annotated regions, making this work a solid proof of concept for automatic tissue classification with machine learning based on HSI data. It is to be considered that automatic semantic scene annotation, however, might still present another challenge. The number of organ recordings is heterogeneous throughout the dataset, which is due to the fact that recordings were done during diverse additional compatible surgical experiments. Moreover, we only considered the median spectrum of one pre-annotated region in one recording, thus ignoring texture information that may further improve organ identification. Notably, excellent classification results could already be achieved despite limiting the neural network input to organ reflectance without texture information. Organs with similar cellular composition such as stomach and jejunum showed similar reflectance spectra, but could still be differentiated well. Misclassifications mainly occurred between bladder and gallbladder, kidney and heart or vena cava and bone.

Besides the investigation of physiological organs, the systematic investigation of pathological states is of likewise importance and need to include tissue ischemia, stasis, inflammation and malignancy. The fact that these unphysiological organ states cannot be purposely induced in patients for ethical reasons necessitates the use of a large animal model with human-like features and known spectral tissue properties and marked the reason for choosing a porcine model for the present study. For proper interpretation of the results of this work, certain limitations inherent to HSI technology have to be taken into account. One limitation is the relatively low temporal resolution of current HSI systems with only one recording every 30 seconds and around seven seconds of recording time each. While more compact and faster devices are under development [57], currently this limitation narrows down possible fields of application. However, it does not undermine the validity of the data presented in this work. In fields of application that require a higher temporal resolution, but not necessarily fine-grained wavelength resolution, multispectral imaging (MSI) offers a solution [4, 58]. These applications might include near video-rate imaging and can most probably be substantially refined when taking insights from HSI research into consideration.

Another limitation of HSI is the generally short and wavelength-dependent penetration depth of light in biological tissue. Increasing penetration depths between 700 and 1,000 nm had to be taken into account when measuring tissue with a thickness of less than several millimeters such as the omentum. Therefore, it was ensured by visual inspection that the omentum was only measured at sites with sufficient thickness. Photoacoustics, as a technology that is able to penetrate more deeply into biological tissue, might help to yield additional information when used complementarily to HSI [59]. Further limitations arise due to the spatial resolution of only 680 × 480 pixels (width × height). Organs with smaller surface areas, e.g. the gallbladder, were harder to annotate than others, since fewer pixels were available. Therefore, the sizes of annotated regions are imbalanced between organs with smaller and larger surface areas. However, small sizes of annotations were successfully compensated by choosing a large number of recordings. Besides the technological limitations, we did not investigate variability resulting from camera variations and the presented tissue identification relied on pre-annotated regions of interest (ROIs). Variability resulting from camera variations and performance on machine learning-based semantic identification will have to be addressed in future studies.

This work is the first to systematically investigate spectral properties and relations of organs within a large cohort of organ classes and individuals. By using a highly standardized approach, we were able to extract the spectral fingerprints for each organ and investigate factors influencing the spectral properties. We were able to provide evidence that the recorded tissues and not the individual animal or the recording conditions were the most influential factor for the electromagnetic spectrum, which is of utmost importance when trying to assess the possible value of HSI for medical applications. This study can be seen as a reference work paving the way for spectral organ evaluation, which requires precise knowledge of spectral characteristics of physiological tissue. Possible future applications based on these results include augmentation of computer-assisted decision-making, intraoperative cognitive assistance systems or even automatization of robotic surgery. It can be expected that our main finding of organ-dependent reflectance patterns will be confirmed in human data. To firmly establish HSI in clinical medicine, a translation of this study to human data will be essential.

## 4 Methods

### 4.1 Animal anaesthesia & surgical procedure

This animal study was approved by the Committee on Animal Experimentation of the regional council of Baden-Württemberg in Karlsruhe, Germany (G-161/18 and G-262/19). All animals used in the experimental laboratory were managed according to German laws for animal use and care, and according to the directives of the European Community Council (2010/63/EU) and ARRIVE guidelines [60]. Data of 46 pigs was included in the analyses.

Experimental animals were operated under general anaesthesia with extensive monitoring including invasive blood pressure measurement. Midline laparotomy was performed to access the abdominal cavity. Ligaments around the liver and the hepato-gastric ligament were dissected and visceral organs mobilized, including the removal of the coverage of the kidneys while carefully sparing vessels. Scissors, electrocautery and bipolar vessel-sealing devices were used. A suprapubic catheter was inserted into the bladder. After surgery, pigs were euthanized with a lethal dose of i.v. potassium chloride solution.

### 4.2 Hyperspectral Imaging

The hyperspectral datacubes were acquired with the TIVITA® Tissue system (Diaspective Vision GmbH, Pepelow, Germany), which is a push-broom scanning imaging system and the first commercially available hyperspectral camera for medicine. It provides a high spectral resolution in the visible as well as near-infrared (NIR) range from 500 nm to 995 nm in 5 nm steps resulting in 100 spectral bands. Its field of view contains 640 × 480 pixels with a spatial resolution of approximately 0.45 mm/pixel (**Figure 5**). The distance of the camera to the specimen is controlled via a red-and-green light targeting system. Six halogen lamps directly integrated into the camera system provide a uniform illumination. Recording takes around seven seconds.

**Figure 5.**
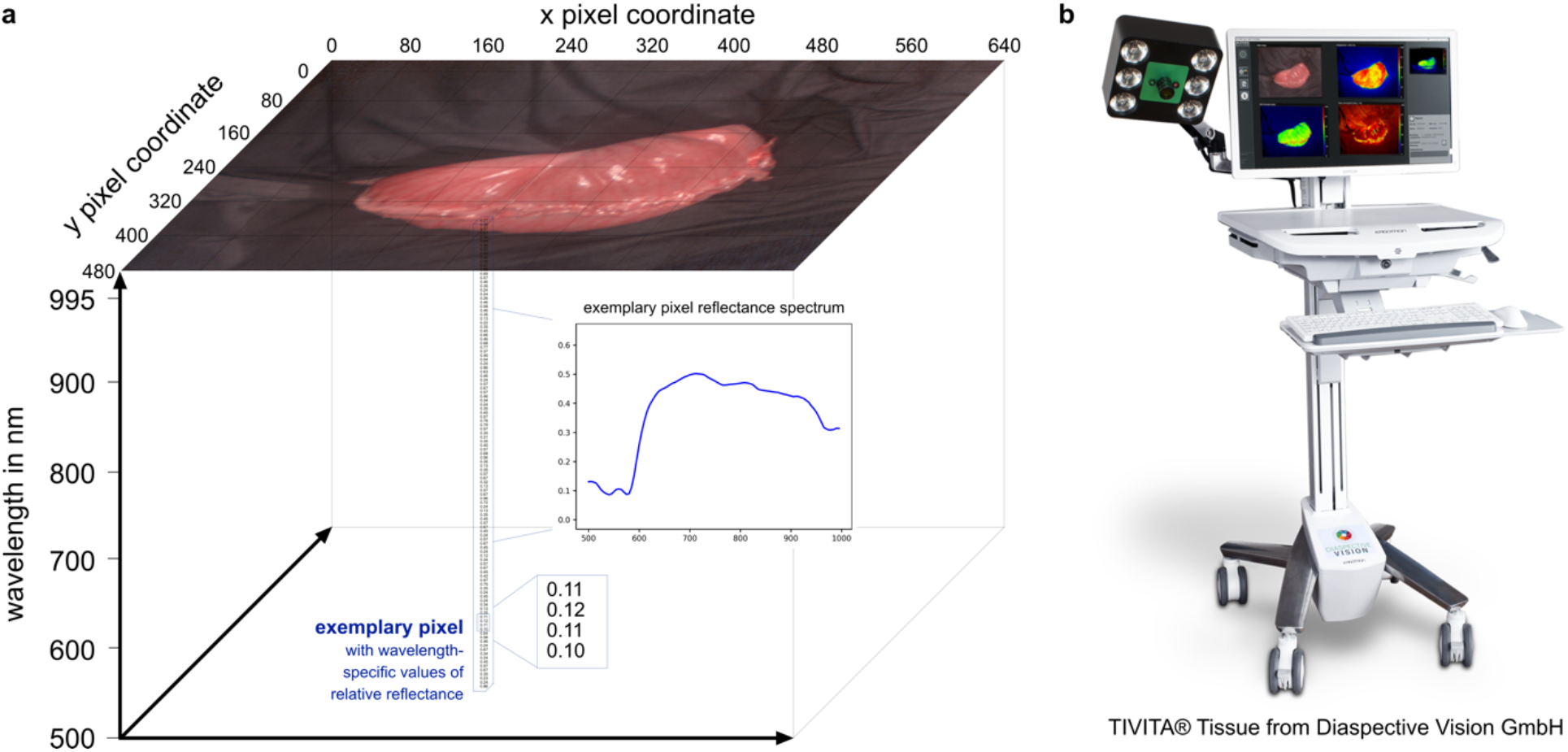
Hyperspectral camera system. **a**, visualization of a three-dimensional hyperspectral datacube with x and y as spatial dimensions and z as hyperspectral dimension. The recorded reflectance information content of one pixel is visualized as an example. **b**, TIVITA® Tissue with kind approval from Diaspective Vision.

### 4.3 Image acquisition, annotation and processing

Images were recorded with a distance of 50 ± 5 cm between camera and organs. In order to prevent distortions of the measured reflectance spectra due to stray light, the tissue recordings were made while lights in the room were switched off and curtains were closed. While the majority of pig recordings was done in a generic approach in order to accurately represent intraoperative reality, recordings for the mixed model analysis were done with a highly standardized protocol for a subset of 11 pigs (8 to 9 pigs per organ) (between P36 and P46 as indicated in **Supplementary Text 1**). This standardized protocol includes recordings of 3 repetitions of exactly the same surgical scene (“repetition” effect) from 3 different angles (“angle” effect) (perpendicular to the tissue surface, 25° from one side and 25° from the opposite side) for 4 different organ positions / situs / situations (“image” effect) resulting in a total of 36 recordings for each of the 20 organs (8 to 9 pigs per organ) in a total of 11 pigs. Recordings for bile fluid were performed when applied and soaked onto 5 stacked surgical compresses, ensuring that there is no influence from the background. For a more extensive overview of the dataset and a schematic recordings protocol for the standardized subset please refer to **Supplementary Figure 1** and **Supplementary Figure 2**.

All of the 9,059 recorded images were sorted into the 20 respective organ folders and manually annotated resulting in 17,777 organ annotations (as several organs could be contained within one image). A precise annotation protocol can be found in **Supplementary Text 5**. Annotations were done by one medical expert and then verified by two other medical experts. In case of improper annotation, the annotation was redone collectively for that specific recording.

For t-SNE and the machine learning analysis, spectral information was previously L1-normalized at pixel-level for increased uniformity. All of the other analyses (including the structured model from **Supplementary Text 2** and **Supplementary Figure 3)** required unprocessed reflectance values from the original datacubes. After annotation, the wavelength-specific annotation-wise median was automatically calculated over every pixel included in the annotation. These median annotation-wise spectra (previously either L1-normalized or not) represented the basic data format that all analyses in this paper were based upon. Calculation of the mean (and SD) integral of the organ reflectance curves of individual animals (**Supplementary Text 3** and **Supplementary Figure 4**) was performed to quantify overall brightness or amount of light that is reflected by the organ in relative units; greater values indicate greater reflectance intensity. Although this quantification of the overall level of the reflectance curve and therefore the area under the curve is influenced by the distance between camera and tissue, the standardization of this distance reduced this influence, rendering this integral a valuable information.

### 4.4 t-SNE

t-distributed Stochastic Neighbor Embedding (t-SNE) [10] is a machine learning method commonly used to reduce the number of dimensions of high-dimensional data and was used to visualize the characteristic reflectance spectra of each pig organ. This non-linear multi-dimensionality reduction tool has already proven valuable for the analysis of HSI and mass spectrometry data [61] and was chosen for visualization as it has shown particular promise for biological samples in the past [62, 63]. The algorithm aims at modelling manifolds of high-dimensional data, and produces low-dimensional embeddings that are optimized for preserving the local neighbourhood structure of the high-dimensional manifold [10]. In comparison to linear methods like PCA [64] and LDA [65], t-SNE preserves more relevant structures of datasets that have non-linear features. For these reasons, t-SNE was used for dimensionality reduction.

Before optimizing the parameters of t-SNE, the dataset was prepared in the following manner: One characteristic reflectance spectrum was obtained for each annotation by calculating the median spectra from the (previously on pixel-level L1-normalized) spectra of all pixels in the annotation. Consequently, each data point represents the reflectance of one organ in one image of one pig. The two-dimensional visualization of the reflectance spectrum of the complete dataset was optimized by performing a random search of the following parameters:

- Parameter 1: The early exaggeration, which controls how tight natural clusters in the original space are in the embedded space and how much space will be between them. 50 random integer values were sampled in the range [5; 100].
- Parameter 2: The learning rate, which is used in the optimization process. 100 random integer values were sampled in the range [10; 1000].
- Parameter 3: The perplexity, which is related to the number of nearest neighbors for each data point to be considered in the optimization. 50 equidistant integer values were sampled uniformly in the range [2; 100].

The early exaggeration was the first parameter optimized by visual inspection of the two-dimensional representation of the dataset. The learning rate was then optimized in the same manner while keeping the early exaggeration constant. Subsequently, the perplexity was optimized by keeping the other two parameters constant. The optimal values for each of the parameters were 34 for the early exaggeration, 92 for the learning rate and 30 for the perplexity.

### 4.5 Linear mixed models

Independent linear mixed models were used for an explained variation analysis in order to evaluate the effect of the influencing factors on changes in the spectrum. The (proportion of) explained variance was obtained using the empirical decomposition of the explained variation in the variance components form of the mixed model [11].

For the first approach, for each wavelength, an independent linear mixed model was fitted with fixed effects for “organ” and “angle” as well as random effects for “pig” and “image”. More precisely, for each wavelength the following model was fitted (suppressing the wavelength index):

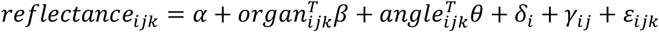

for repetition *k*=1,…,3 of image *j*=1,…,n_i_ of pig *i*=1,…, 11 (with n_i_ the number of images of pig *i* ranging from 84 to 228 and 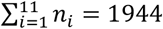 + is an intercept, 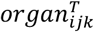 is a row vector of length 19 indicating the organ of observation *ijk* (with arbitrary reference category “stomach”) and *β* is a vector of corresponding fixed organ effects. Similarly, *θ* are fixed effects for angle (“25° from one side” and “25° from the opposite side” for reference category “perpendicular to the tissue surface”). 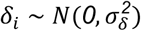 and 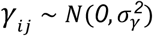 are random pig and image effects, respectively, assumed to be independently normal distributed with between pig variation 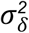 and between image variation 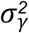 Residuals 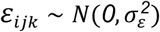 capture the variability between repeated recordings of the same image.

The proportion of variability in reflectance explained by each factor was derived as in [11]. “Repetition” depicts the residual variability, which is here the within image variability (i.e. across replications). 95 % pointwise confidence intervals based on parametric bootstrapping with 500 replications indicate the uncertainty in estimates.

For the second approach with stratification by organ, independent linear mixed models were fitted for each organ and wavelength with fixed effects for “angle” as well as a random effect for “pig” and “image”, i.e for each organ and wavelength the same model as given above was fitted excluding covariate “organ”. The explained variation of each factor was depicted [11]. “Repetition” depicts the residual variation, which is here the within image variability (i.e. across replications). 95 % pointwise confidence intervals based on parametric bootstrapping with 500 replications indicate the uncertainty in estimates. Curves were linearly interpolated if model fit was singular. All linear mixed model analyses were based on image-wise organ-specific median reflectance spectra that were obtained by calculating the median spectrum of all pixel spectra within one annotation.

### 4.6 Machine Learning

Prior to training our deep learning network, we systematically split the dataset comprising 46 pigs (9,059 images with 17,777 annotations) into a training dataset consisting of 38 pigs (3,766 images with 7,882 annotations) and a disjoint test set consisting of 8 pigs (5,293 images with 9,895 annotations) as indicated in **Supplementary Figure 1**. These 8 test pigs were randomly selected from the 11 standardized pigs (P36-P46) with the only criterion that every organ class is represented by at least one standardized pig in the test as well as in the training dataset. This criterion could not be fulfilled anymore when selecting more than 8 standardized pigs.

The hold-out test set was used only after the network architecture and all hyperparameters had been fixed. Leave-one-pig-out cross-validation was performed on the training dataset and the predictions on the left-out pig were aggregated for all 38 folds (46 minus 8) to yield the validation accuracy. The hyperparameters of the neural network were optimized in an extensive grid search such that the validation accuracy was maximized. Once the optimal hyperparameters were determined, we evaluated the classification performance on the hold-out test set by ensembling the predictions from all 38 networks (one for each fold) via computing the mean logits vector (the input values to the softmax function, see below) followed by the argmax operation to retrieve the final label for each annotation.

The deep learning-based classification was performed on the median spectra computed from the L1-normalized spectra of all pixels in the annotation masks resulting in 100-dimensional input feature vectors.

The deep learning architecture was composed of 3 convolutional layers (64 filters in the first, 32 in the second, and 16 in the third layer) followed by 2 fully connected layers (100 neurons in the first and 50 in the second layer). The activations of all five layers were batch normalized and a final linear layer was used to calculate the class logits. Each of the convolutional layers convolved the spectral domain with a kernel size of 5 and was followed by an average pooling layer with a kernel size of 2. The two fully connected layers zeroed out their activations with a dropout probability of ã. All non-linear layers used the Exponential Linear Unit (ELU) [66] as activation function.

We chose this architecture as it provides a simple yet effective way to analyze the spectral information. The convolution operation acts on the local structure of the spectra and we used a relatively small kernel size and stacked 3 layers to increase the receptive field while being computationally efficient [67]. The two fully connected layers make a final decision based on the global context. The advantage of this approach is that it combines local and global information aggregation while still being computationally efficient since the entire network only uses 34,300 trainable weights. The softmax function was used to provide the *a posteriori* probability for each class. We used the Adam optimizer (β_1_= 0.9, β_2_= 0.999) [68] with an exponential learning rate decay (decay rate of γ and initial learning rate of *η*) and the multiclass cross-entropy loss function. In order to meet class imbalances, we included an optional weight of the loss function according to the number of training images per class and sampled instances for the batches either randomly or oversampled such that each organ class had the same probability of being sampled. Both design choices were investigated in the hyperparameter grid search.

We trained 10,000,000 samples per epoch for 10 epochs with a batch size of *N*. In an extensive grid search, we determined the best-performing hyperparameters: dropout probability *p*^*^ = 0.2 *p* ∈ {0.1, 0.2}), learning rate *η* ^*^ = 0.0001 (*η* ∈ {0.001, 0.0001}), decay rate *γ* ^*^ = 0.9 (*γ*; ∈ {0.75, 0.9, 1.0}),batch size *N*^*^ =20,000 (*N* ∈ {20,000, 40,000}), a weighted loss function and no oversampling.

## Abbreviations

HIS: hyperspectral imaging
MSI: multispectral imaging
NASA: National Aeronautics and Space Administration
ROI: region of interest
SD: standard deviation
t-SNE: t-distributed Stochastic Neighbor Embedding

## Acknowledgements

The authors thank Minu Tizabi (CAMI, DKFZ) for proofreading the manuscript and gratefully acknowledge the data storage service SDS@hd supported by the Ministry of Science, Research and the Arts Baden-Württemberg (MWK) and the German Research Foundation (DFG) through grant INST 35/1314-1 FUGG and INST 35/1503-1 FUGG. The present contribution is supported by the Helmholtz Association under the joint research school HIDSS4Health (Helmholtz Information and Data Science School for Health). This project received funding from the European Research Council (ERC) under the European Union’s Horizon 2020 research and innovation program (NEURAL SPICING, grant agreement No. [101002198]) as well as from the Willi Robert Pitzer foundation, from the Heidelberg Foundation for Surgery and from the RISE program by the German Academic Exchange Service (DAAD).

## Author contributions

ASF, FN and BPMS had the original idea for the project. ASF and FN initiated the project. ASF, FN and CMH performed the initial review of existing literature and the planning. ASF, CMH, KK, BÖ, MD, BPMS and FN performed the surgeries. ASF, IC, BÖ and JO developed the Python codes for data organization and annotation. ASF, GS and SK annotated data. ASF, GS, BÖ, JO, SK, SS, LA, JS, LMH, TJA and MW analyzed and interpreted data. LMH, TJA, LA, SS, JS, MW, NS, AKS and ASF developed and implemented the statistical and machine learning-based data analysis strategy. LA provided the t-SNE analysis and the relevant manuscript passages. MW, NS and AKS provided the mixed model analysis, the structured model and the relevant manuscript passages. SS and JS provided the machine learning-based classification and the relevant manuscript passages. FN, LMH, HGK, KMH, KS and BPMS provided expert knowledge throughout the project. ASF, BÖ and FN wrote the manuscript. SS, JS, LA, MW, BPMS, HGK, KMH and LMH revised the manuscript. All authors have read and approved the final manuscript.

## Statement about competing interests

Authors state no conflict of interest.

## References

1. Mascagni, P., et al., New intraoperative imaging technologies: Innovating the surgeon’s eye toward surgical precision. Journal of Surgical Oncology, 2018. 118(2): p. 265–282.

2. Prasad, S. and J. Chanussot, Hyperspectral Image Analysis - Advances in Machine Learning and Signal Processing. Advances in Computer Vision and Pattern Recognition. 2020: Springer.

3. Ayala, L.A., et al. Live Monitoring of Haemodynamic Changes with Multispectral Image Analysis. 2019. Cham: Springer International Publishing.

4. Wirkert, S.J., et al. Physiological Parameter Estimation from Multispectral Images Unleashed. 2017. Cham: Springer International Publishing.

5. Barberio, M., et al., Quantitative fluorescence angiography versus hyperspectral imaging to assess bowel ischemia: A comparative study in enhanced reality. Surgery, 2020.

6. Nickel*, F. & Studier-Fischer*, A., et al., Optimization of anastomotic technique and gastric conduit perfusion with hyperspectral imaging in an experimental model for minimally invasive esophagectomy. bioRxiv, 2021: p. 2021.10.03.462901.

7. Dietrich*, M. & Seidlitz*, S., et al. Machine learning-based analysis of hyperspectral images for automated sepsis diagnosis. arXiv 2021 20.08.2021]; Available from: https://arxiv.org/abs/2106.08445v1.

8. Wu, I.C., et al., Early identification of esophageal squamous neoplasm by hyperspectral endoscopic imaging. Sci Rep, 2018. 8(1): p. 13797.

9. Clancy, N.T., et al., Surgical spectral imaging. Medical Image Analysis, 2020. 63: p. 101699.

10. van der Maaten, L.J.P. and G.E. Hinton, Visualizing High-Dimensional Data Using t-SNE. Journal of Machine Learning Research, 2008. 9(86): p. 2579™2605.

11. Schreck, N., Empirical decomposition of the explained variation in the variance components form of the mixed model. bioRxiv, 2019: p. 2019.12.28.890061.

12. Thunell, S., Porphyrins, porphyrin metabolism and porphyrias. I. Update. Scand J Clin Lab Invest, 2000. 60(7): p. 509–40.

13. Wilson, M.T. and B.J. Reeder, Oxygen-binding haem proteins. Exp Physiol, 2008. 93(1): p. 128–32.

14. Jacques, S.L., Optical properties of biological tissues: a review. Physics in Medicine and Biology, 2013. 58(11): p. R37–R61.

15. Lu, G. and B. Fei, Medical hyperspectral imaging: a review. Journal of Biomedical Optics, 2014. 19(1): p. 010901.

16. Afromowitz, M.A., et al., Multispectral imaging of burn wounds: a new clinical instrument for evaluating burn depth. IEEE Transactions on Biomedical Engineering, 1988. 35(10): p. 842–850.

17. Ferris, D.G., et al., Multimodal Hyperspectral Imaging for the Noninvasive Diagnosis of Cervical Neoplasia. Journal of Lower Genital Tract Disease, 2001. 5(2): p. 65–72.

18. Shah, S.A., et al., Cutaneous Wound Analysis Using Hyperspectral Imaging. BioTechniques, 2003. 34(2): p. 408–413.

19. Bambery, K.R., et al., Fourier Transform Infrared Imaging and Unsupervised Hierarchical Clustering Applied to Cervical Biopsies. Australian Journal of Chemistry, 2004. 57(12): p. 1139–1143.

20. Akbari, H., et al., Detection and Analysis of the Intestinal Ischemia Using Visible and Invisible Hyperspectral Imaging. IEEE Transactions on Biomedical Engineering, 2010. 57(8): p. 2011–2017.

21. Akbari, H., et al., Hyperspectral imaging and quantitative analysis for prostate cancer detection. Journal of Biomedical Optics, 2012. 17(7): p. 076005.

22. Mitra, K., et al., Indocyanine-green-loaded microballoons for biliary imaging in cholecystectomy. J Biomed Opt, 2012. 17(11): p. 116025.

23. Rosas, J.G. and M. Blanco, A criterion for assessing homogeneity distribution in hyperspectral images. Part 2: Application of homogeneity indices to solid pharmaceutical dosage forms. Journal of Pharmaceutical and Biomedical Analysis, 2012. 70: p. 691–699.

24. Li, Q., et al. Nerve fibers identification based on molecular hyperspectral imaging technology. in 2012 IEEE International Conference on Computer Science and Automation Engineering (CSAE). 2012.

25. Kumar, S., et al., Change in the microenvironment of breast cancer studied by FTIR imaging. Analyst, 2013. 138(14): p. 4058–65.

26. Grambow, E., et al., Hyperspectral imaging for monitoring of perfusion failure upon microvascular anastomosis in the rat hind limb. Microvasc Res, 2018. 116: p. 64–70.

27. Barberio, M., et al., HYPerspectral Enhanced Reality (HYPER): a physiology-based surgical guidance tool. Surgical Endoscopy, 2020. 34(4): p. 1736–1744.

28. Felli, E., et al. Hyperspectral imaging of pig liver ischemia: a proof of concept. 2019 20.08.2021]; Available from: https://www.airitilibrary.com/Publication/alDetailedMesh?docid=15610497-201912-201912180004-201912180004-117-121.

29. Tetschke, F., et al., Hyperspectral imaging for monitoring oxygen saturation levels during normothermic kidney perfusion. J. Sens. Sens. Syst., 2016. 5(2): p. 313–318.

30. Holmer, A., et al. Bildgebende chemische Analyse und die Anwendung in der medizinischen Perfusions-Forschung. in AUTOMED. 2016. Wismar.

31. Landro, M.D., et al. Hyperspectral imaging for thermal effect monitoring in in vivo liver during laser ablation. in 2019 41st Annual International Conference of the IEEE Engineering in Medicine and Biology Society (EMBC). 2019.

32. Gockel, I., et al., Möglichkeiten und Perspektiven der Hyperspektralbildgebung in der Viszeralchirurgie. Der Chirurg, 2020. 91(2): p. 150–159.

33. Goetze, E., et al., Digitalisierung und Ansätze künstlicher Intelligenz in der mikrovaskulär-rekonstruktiven Gesichtschirurgie. Der Chirurg, 2020. 91(3): p. 216–221.

34. Maier-Hein, L., et al., Intraoperative Bildgebung und Visualisierung. Der Onkologe, 2020. 26(1): p. 31–43.

35. Holmer, A., et al., Oxygenation and perfusion monitoring with a hyperspectral camera system for chemical based tissue analysis of skin and organs. Physiol Meas, 2016. 37(11): p. 2064–2078.

36. Markgraf, W., et al., Algorithms for mapping kidney tissue oxygenation during normothermic machine perfusion using hyperspectral imaging. Biomedical Engineering / Biomedizinische Technik, 2018. 63(5): p. 557.

37. Barberio, M., et al., Hyperspectral based discrimination of thyroid and parathyroid during surgery. 2018. 4(1): p. 399.

38. Daeschlein, G., et al., Hyperspectral imaging as a novel diagnostic tool in microcirculation of wounds. Clin Hemorheol Microcirc, 2017. 67(3-4): p. 467–474.

39. Grambow, E., et al., Evaluation of peripheral artery disease with the TIVITA(R) Tissue hyperspectral imaging camera system. Clin Hemorheol Microcirc, 2019. 73(1): p. 3–17.

40. Herrmann, B.H. and C. Hornberger, Monte-Carlo Simulation of Light Tissue Interaction in Medical Hyperspectral Imaging Applications. Current Directions in Biomedical Engineering, 2018. 4(1): p. 275–278.

41. Jansen-Winkeln, B., et al., Bestimmung der idealen Anastomosenposition durch hyperspectrale Bildgebung. Z Gastroenterol, 2019. 57(09): p. KV 93.

42. Jansen-Winkeln, B., et al., Determination of the transection margin during colorectal resection with hyperspectral imaging (HSI). International Journal of Colorectal Disease, 2019. 34(4): p. 731–739.

43. Jansen-Winkeln, B., et al., Handnaht v. Stapler-Anastomose – Hyperspektralbetrachtung der Perfusion. Z Gastroenterol, 2019. 57(09): p. KV 91.

44. Köhler, H., et al., Hyperspectral imaging as a new optical method for the measurement of gastric conduit perfusion. Diseases of the Esophagus, 2019. 32(10): p. 1–1.

45. Köhler, H., et al., Evaluation of hyperspectral imaging (HSI) for the measurement of ischemic conditioning effects of the gastric conduit during esophagectomy. Surgical Endoscopy, 2019. 33(11): p. 3775–3782.

46. Langner, I., et al., Hyperspektralimaging demonstriert mikrozirkulatorische Effekte postoperativer Ergotherapie bei Patienten mit Morbus Dupuytren. Handchir Mikrochir plast Chir, 2019. 51(03): p. 171–176.

47. Maktabi, M., et al., Tissue classification of oncologic esophageal resectates based on hyperspectral data. International Journal of Computer Assisted Radiology and Surgery, 2019. 14(10): p. 1651–1661.

48. Marotz, J., et al., Extended Perfusion Parameter Estimation from Hyperspectral Imaging Data for Bedside Diagnostic in Medicine. Molecules (Basel, Switzerland), 2019. 24(22): p. 4164.

49. Marotz, J., et al., First results of a new hyperspectral camera system for chemical based wound analysis. Wound Medicine, 2015. 10-11: p. 17–22.

50. Mohammed, R.A.A., et al., Detecting Signatures in Hyperspectral Image Data of Wounds: A Compound Model of Self-Organizing Map and Least Square Fitting. Current Directions in Biomedical Engineering, 2018. 4(1): p. 419–422.

51. Sucher, R., et al., Hyperspectral Imaging (HSI) in anatomic left liver resection. International Journal of Surgery Case Reports, 2019. 62: p. 108–111.

52. Zimmermann, P., et al., Analysis of tissue oxygenation in chronic leg ulcers by combination of a multi-spectral camera and a hyper-spectral probe. Georgian Med News, 2017(270): p. 75–81.

53. Maktabi, M., et al., Tissue classification of oncologic esophageal resectates based on hyperspectral data. Int J Comput Assist Radiol Surg, 2019.

54. Filatova, S., I. Shcherbakov, and V. Tsvetkov, Optical properties of animal tissues in the wavelength range from 350 to 2600 nm. Journal of Biomedical Optics, 2017. 22(3): p. 035009.

55. Zhang, Y., et al., Tissue classification for laparoscopic image understanding based on multispectral texture analysis. J Med Imaging (Bellingham), 2017. 4(1): p. 015001.

56. Moccia, S., et al., Uncertainty-Aware Organ Classification for Surgical Data Science Applications in Laparoscopy. IEEE Trans Biomed Eng, 2018. 65(11): p. 2649–2659.

57. Ayala, L., et al. Video-rate multispectral imaging in laparoscopic surgery: First-in-human application. 2105.13901, 2021.

58. Wirkert, S.J., et al., Robust near real-time estimation of physiological parameters from megapixel multispectral images with inverse Monte Carlo and random forest regression. Int J Comput Assist Radiol Surg, 2016. 11(6): p. 909–17.

59. Gröhl, J., et al. Semantic segmentation of multispectral photoacoustic images using deep learning. 2105.09624, 2021.

60. Kilkenny, C., et al., Animal research: reporting in vivo experiments: the ARRIVE guidelines. Br J Pharmacol, 2010. 160(7): p. 1577–9.

61. Gardner, W., et al., Understanding mass spectrometry images: complexity to clarity with machine learning. Biopolymers, 2020. n/a(n/a): p. e23400.

62. Kobak, D. and P. Berens, The art of using t-SNE for single-cell transcriptomics. Nature Communications, 2019. 10(1): p. 5416.

63. Cieslak, M.C., et al., t-Distributed Stochastic Neighbor Embedding (t-SNE): A tool for eco-physiological transcriptomic analysis. Marine Genomics, 2020. 51: p. 100723.

64. Tipping, M.E. and C.M. Bishop, Probabilistic Principal Component Analysis. Journal of the Royal Statistical Society: Series B (Statistical Methodology), 1999. 61(3): p. 611–622.

65. Hastie, T., R. Tibshirani, and J. Friedman, The Elements of Statistical Learning. 2 ed. Springer Series in Statistics. 2008: Springer. 106–119.

66. Clevert, D.-A., T. Unterthiner, and S. Hochreiter. Fast and Accurate Deep Network Learning by Exponential Linear Units (ELUs). 2016 20.08.2021]; Identifier: 1511.07289]. Available from: http://arxiv.org/abs/1511.07289.

67. Szegedy, C., et al. Rethinking the Inception Architecture for Computer Vision. in 2016 IEEE Conference on Computer Vision and Pattern Recognition (CVPR). 2016.

68. Kingma, D.P. and J. Ba. Adam: A method for stochastic optimization. arXiv preprint 1412.6980 2014 20.08.2021]; Available from: https://arxiv.org/abs/1412.6980.

69. Wood, S.N., Fast stable restricted maximum likelihood and marginal likelihood estimation of semiparametric generalized linear models. Journal of the Royal Statistical Society: Series B (Statistical Methodology), 2011. 73(1): p. 3–36.

70. Pedersen, E.J., et al., Hierarchical generalized additive models in ecology: an introduction with mgcv. PeerJ 7:e6876, 2019.

