## Supplement for "Spectral organ fingerprints for intraoperative tissue classification with hyperspectral imaging"

### Supplementary Text 1: Data overview and standardized recordings protocol

The complete dataset for this study consists of 9,059 images of 46 pigs with a total of 17,777 annotations for 20 different organ classes. The exact data distribution can be seen in **Supplementary Figure 1**. It is important to note that one recording can have multiple organs in its field of view and therefore can have multiple organs annotated. Hence, the total number of annotations across all organs is greater than the total number of images in the data set. Within one organ, the total number of recordings equals the total number of annotations. Every image has at least one organ annotation. The number of recordings per animal is heterogeneous since the number of “standardized” images taken for pigs P36 to P46 is generally higher than the number of images taken when imitating intraoperative reality. Furthermore, the distribution of number of images per organ is heterogeneous since some organs naturally occur more often in the field of view of other organs, e.g. during recordings of the gallbladder, the liver is always in the same field of view but for liver recordings, the gallbladder is not always apparent. However, all analyses were performed in a manner independent of neighboring structures. The linear mixed models analysis was done on the recordings marked with scratched boxes in **Supplementary Figure 1**. Testing of the machine learning algorithm was performed on all images of the 8 animals marked with asterisks.

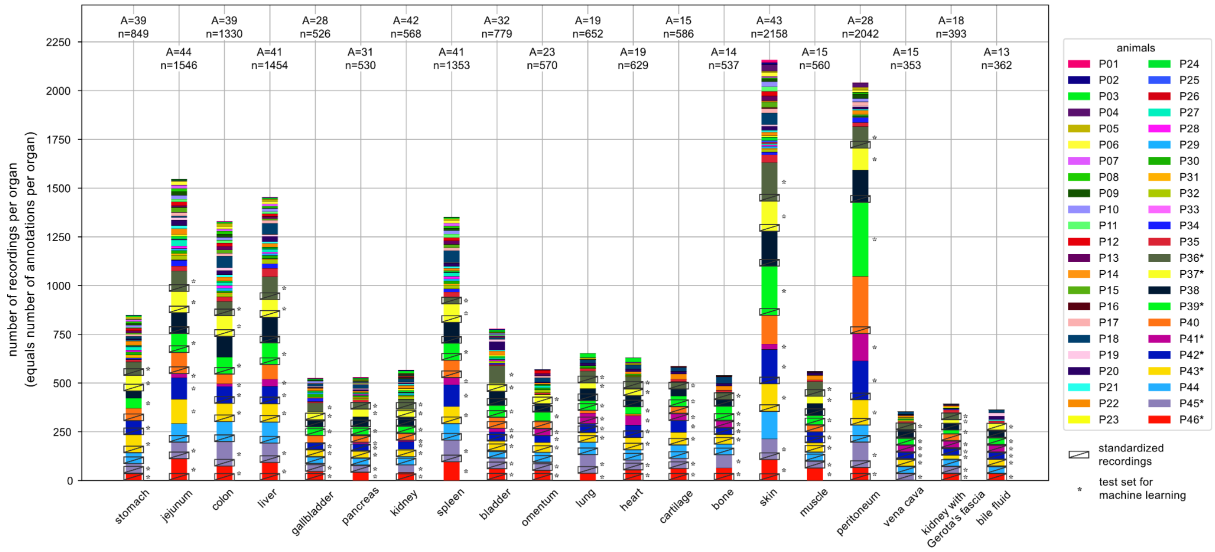


**Supplementary Figure 1 | Database visualization.** One color corresponds to one pig with P01 at the top and P46 at the bottom of the bar chart. The scratched box indicates the recordings from the 11 standardized pig measurements that were used for the linear mixed models analysis with 36 recordings per box and 8 boxes per organ. The asterisk indicates the 8 pigs that were used as a test set and for reporting accuracy of the machine learning algorithm.

The standardized images were recorded with a specific protocol, which can be seen illustrated in **Supplementary Figure 2** including 3 repetitions in 3 angles in 4 situs of each organ (for 8 pigs per organ) in a total of 11 pigs.

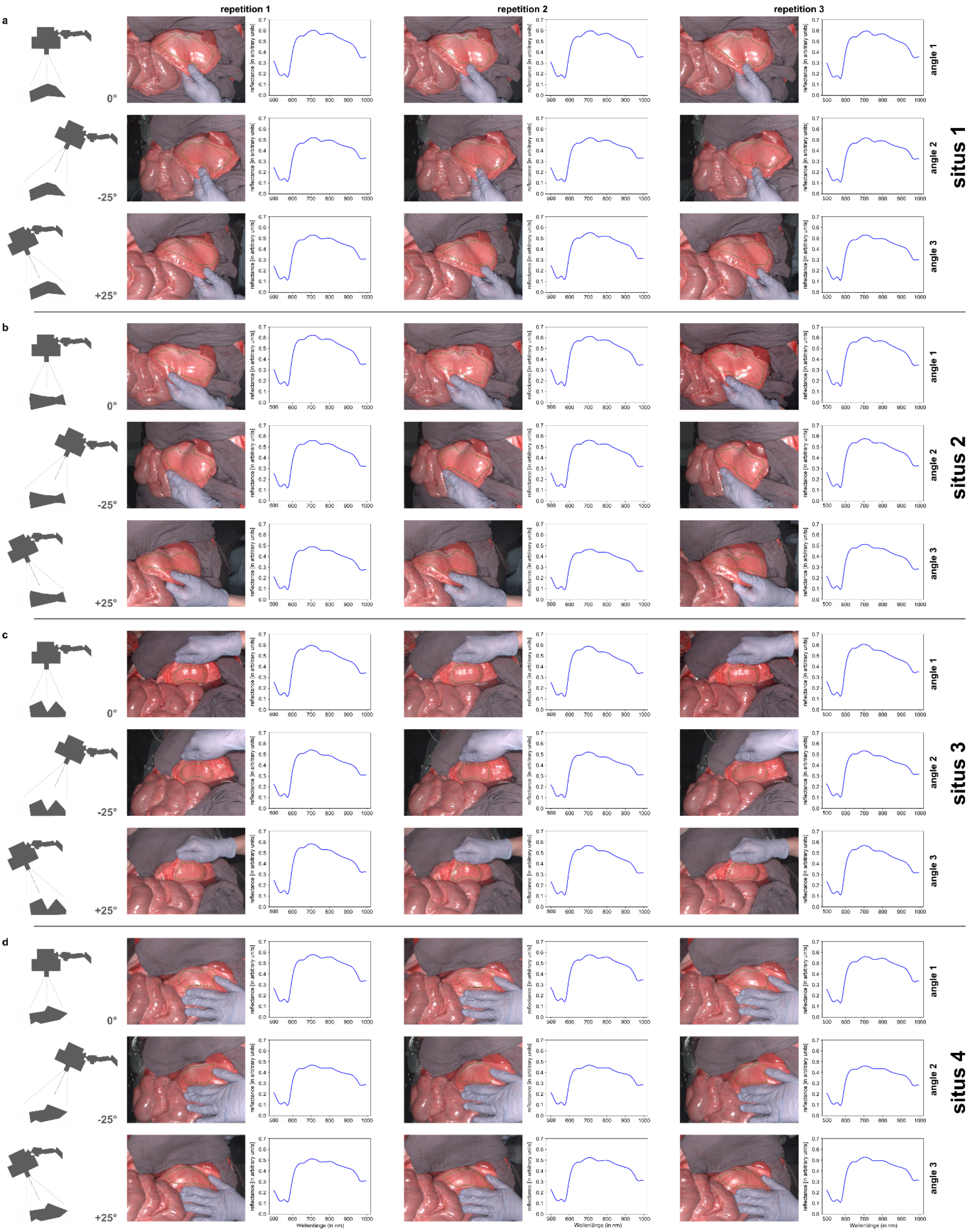


**Supplementary Figure 2 | Schematic recordings protocol for the standardized dataset.** The schematic visualization depicts the 36 recordings that were taken per organ in 8 animals. These include 3 repetitions in 3 angles in 4 situs. A stomach recording is used as an example together with the extracted reflectance spectrum of the corresponding annotation.

**Supplementary Text 2: Structured model analysis**

In order to enhance our understanding of the single influencing components that make up the recorded spectra, the complete dataset was systematically deconstructed and consecutively reconstructed through a structured model [69]. For this, an additive model was fitted regressing reflectance (assuming a Gaussian distribution) on a smooth overall mean curve along wavelength plus smooth deviations from this curve by organs plus a smooth random effect for “pig” and a random intercept for “image” as well as a fixed effect for “angle” was calculated (**Supplementary Figure 3**). P-splines with 25 knots were used for estimation of smooth terms, except for smooth random pig effects where 10 knots were used. First-order penalty was used for smooth organ-specific deviations and quadratic penalties otherwise [70]. The high similarity between raw data and its prediction indicates how the entire dataset has clearly nameable components that contribute to the final shape of reflectance.

The structured model provides a different view on hyperspectral tissue characterizations of organs shown in **Figure 1** by decomposing the wavelength-reflectance curves into an overall mean curve shared by all organs (a) and organ-specific deviations from this overall mean curve (d) as well as smooth pig-specific deviations from the overall mean modelled as random effects (b) (**Supplementary Figure 3**). The organ-specific deviations in (d) can be further decomposed into a general mean level shift from the overall mean constant across wavelengths (c top) and differences in the shape of the organ-specific curves (c bottom). Panel (d) illustrates that reflectance curves were relatively constant for each organ across wavelengths above a wavelength of approximately 600, but on distinct levels. Panel (c top) shows which organs shared a similar mean reflectance level across all wavelengths. Bile fluid was intuitively shown to have a very different reflectance profile than the solid organs most strikingly visible in (c bottom). Panel (b) illustrates that after the organ effect was controlled for, the pig-specific variation of reflectance was relatively small across wavelengths (with curves between approximately -0.04 and 0.06) as compared to differences between organs with organ-specific deviations from the overall mean between +/- 0.35. Angle effects were small with -0.027 (SE 0.003) for 25° from body-left and 0.041 (SE 0.003) for category. 95 % of predicted image-specific deviations (image random effects) range between +/- 0.1, reflecting the same finding as in the mixed model analysis that image is the second strongest effect and potentially more important than pig. Panel (e) illustrates that the model is suitable to predict reflectance relationships.

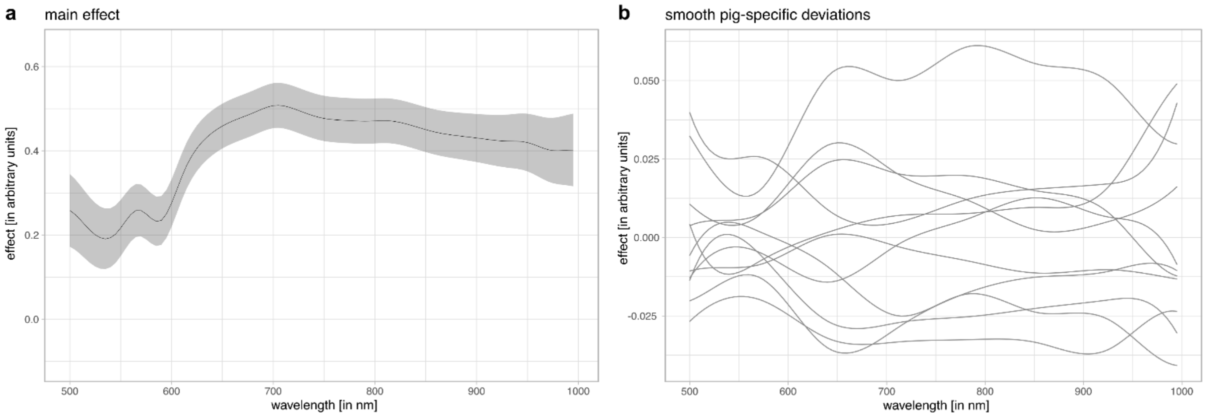

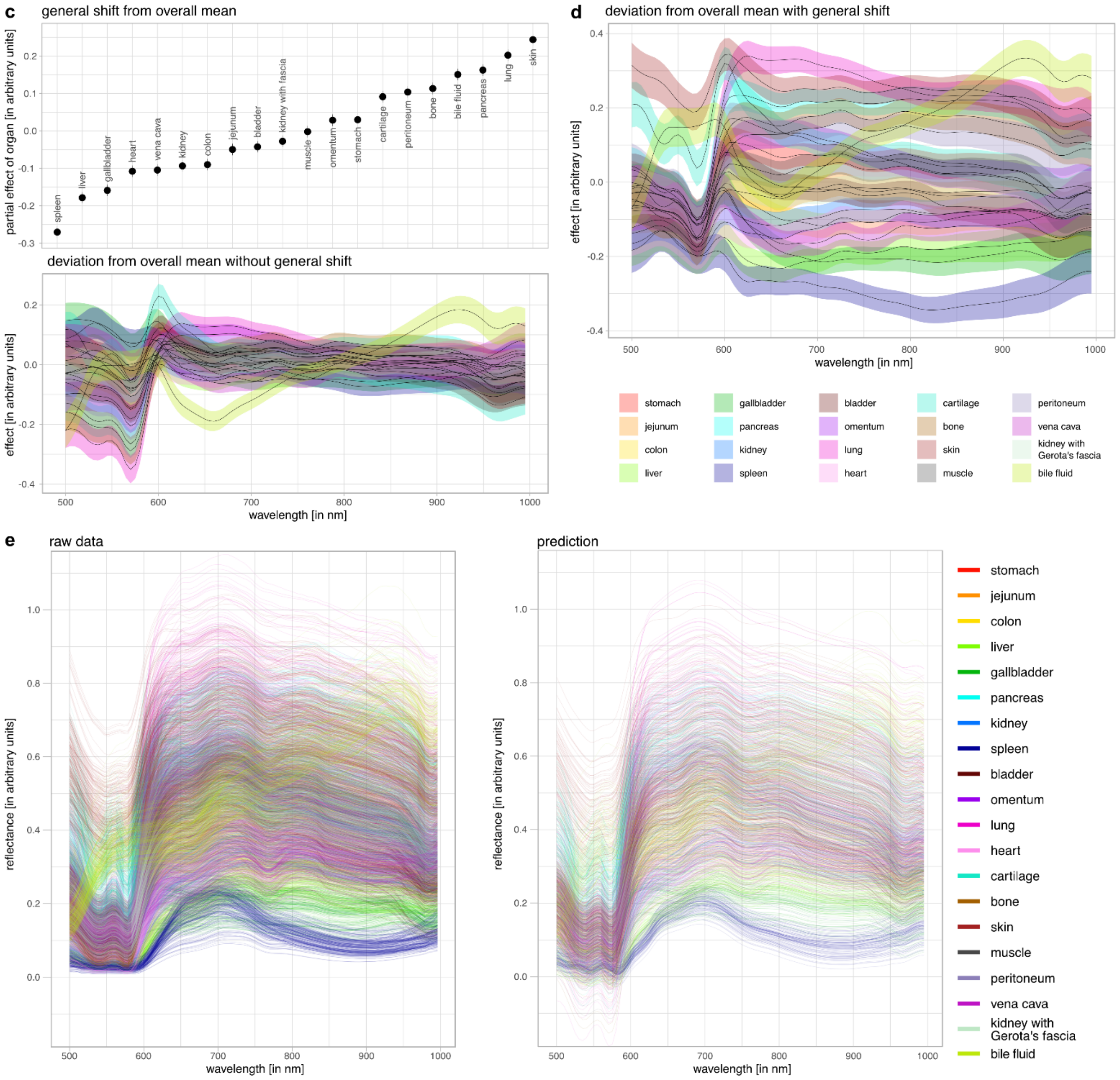


**Supplementary Figure 3 | Structured model. a**, Overall effect: displays the overall mean wavelength-reflectance relationship shared by all. **b**, pig random smooth effect independent of image and organ and overall mean. **c**, organ level shift (displays the level shift from the overall mean reflectance relationship for each organ) and deviation from overall mean (displays the deviation from the overall mean reflectance relationship for each organ; excluding level shift). **d**, organ-specific deviation from overall mean including level shift. Angle and image effects were not visualized as they can be sufficiently described in the text. **a-d**: 95 % pointwise confidence bands i.e. asymptotic pointwise confidence intervals based on normal distribution. **e**, raw data (on annotation level with one curve for every annotation of every organ in every pig) and prediction (on image level with one curve for every image of every organ in every pig) from the model; note that the model does not respect that reflectance is effectively bounded by below zero.

**Supplementary Text 3: Hyperspectral tissue atlas**

Representative color-coded index pictures of physiological organs are depicted together with their respective reflectance and absorption graphs as well as their first and second derivative in order to provide a comprehensive spectral tissue atlas of physiological organs and to visualize organ-characteristic features (**Supplementary Figure 4**). All these different graphs provide an enhanced overview and understanding for organ differentiation opportunities in that even organs with only subtle differences in reflectance can much more easily be differentiated subjectively by inspecting larger differences in the derivative graphs. Height of the reflectance curve is quantified using the integral and can be seen as overall reflectance intensity or “brightness”. Maximum values for “brightness” were obtained for lung (56.1; SD 8.1); minimum values for spleen (8.5; 1.6).

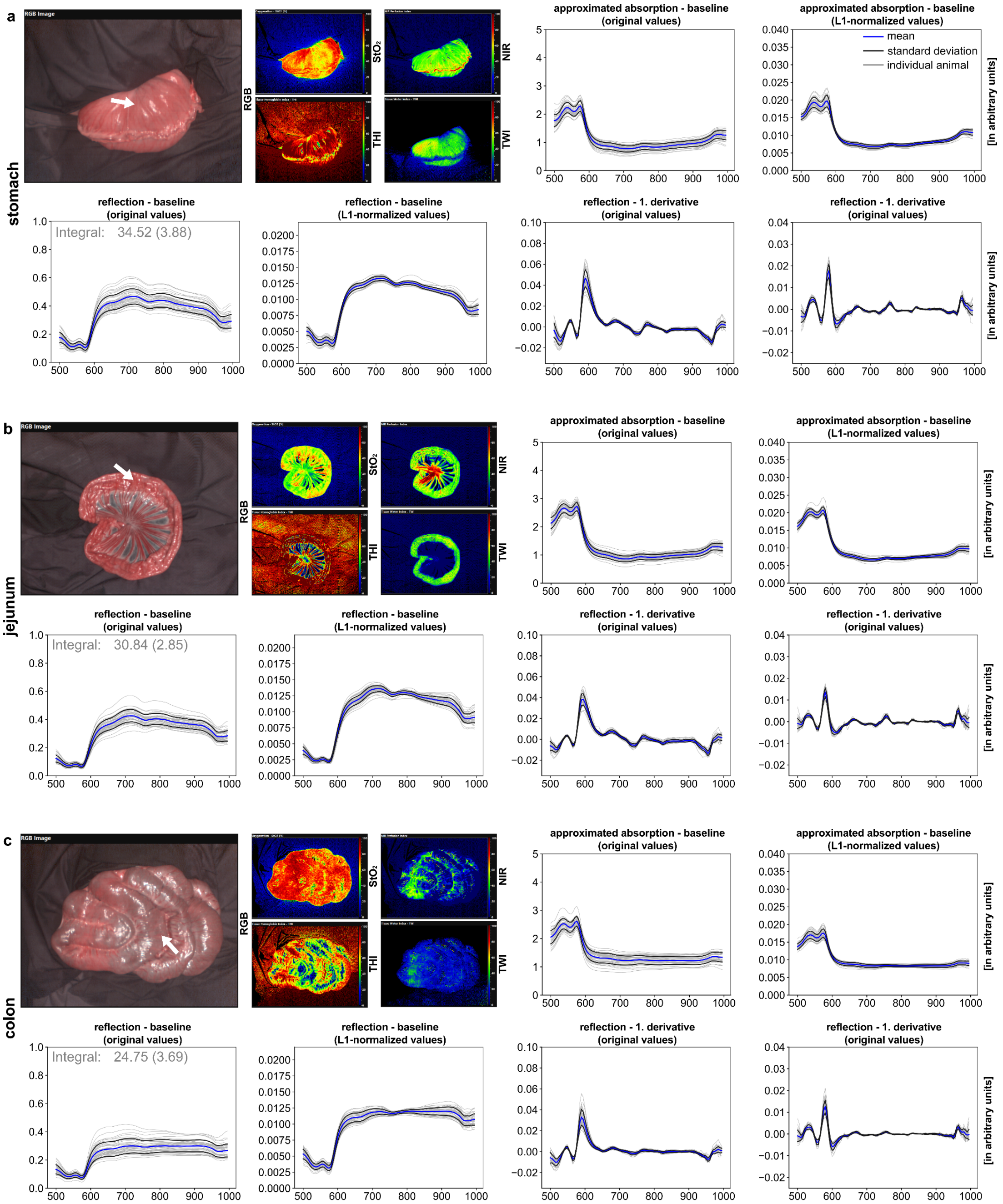

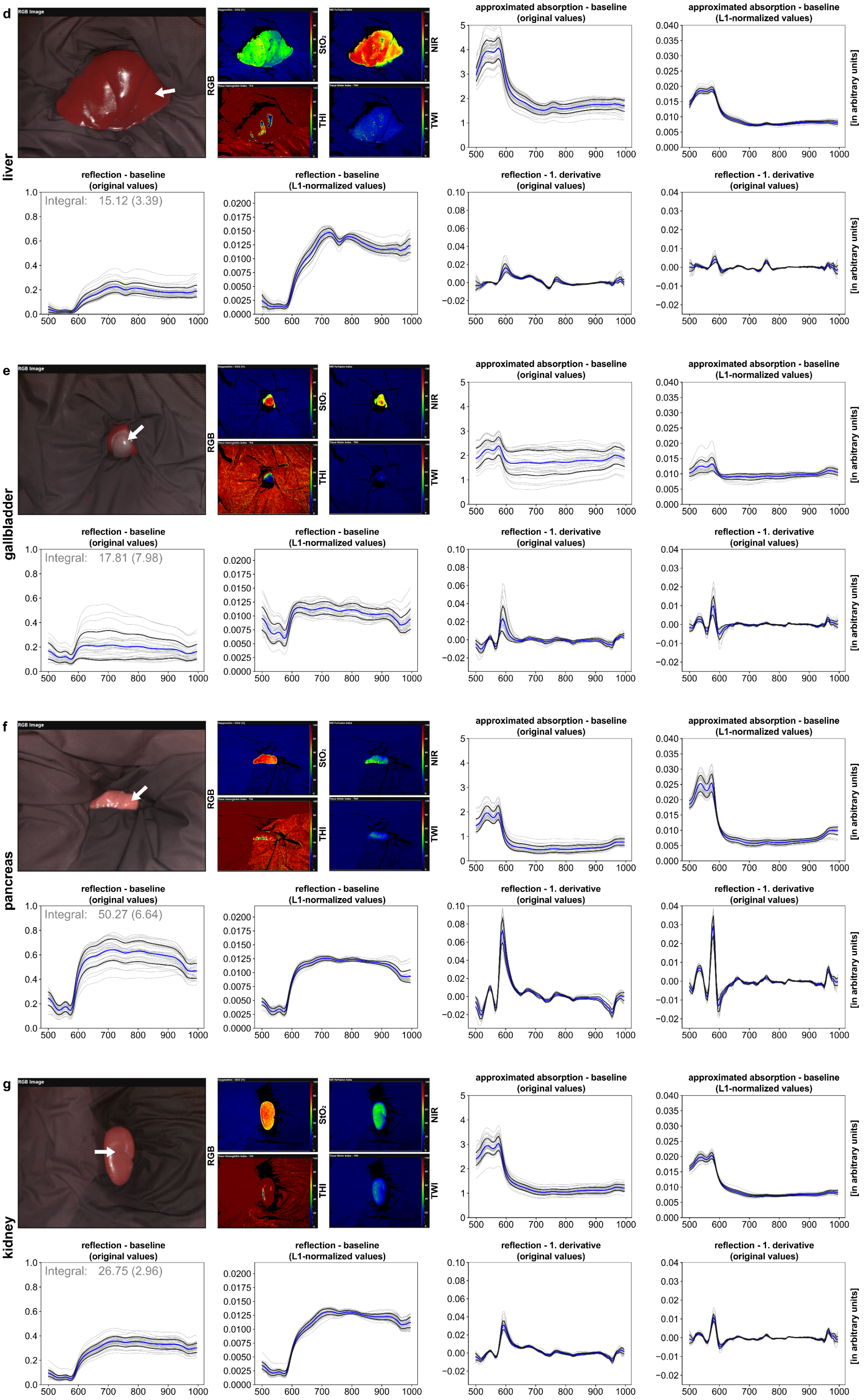
d
liver
RGB Image
RGB
StO₂
NIR
THI
TWI
approximated absorption - baseline (original values)
approximated absorption - baseline (L1-normalized values)
[in arbitrary units]
reflection - baseline (original values)
Integral: 15.12 (3.39)
reflection - baseline (L1-normalized values)
reflection - 1. derivative (original values)
reflection - 1. derivative (original values)
e
gallbladder
Integral: 17.81 (7.98)
f
pancreas
Integral: 50.27 (6.64)
g
kidney
Integral: 26.75 (2.96)

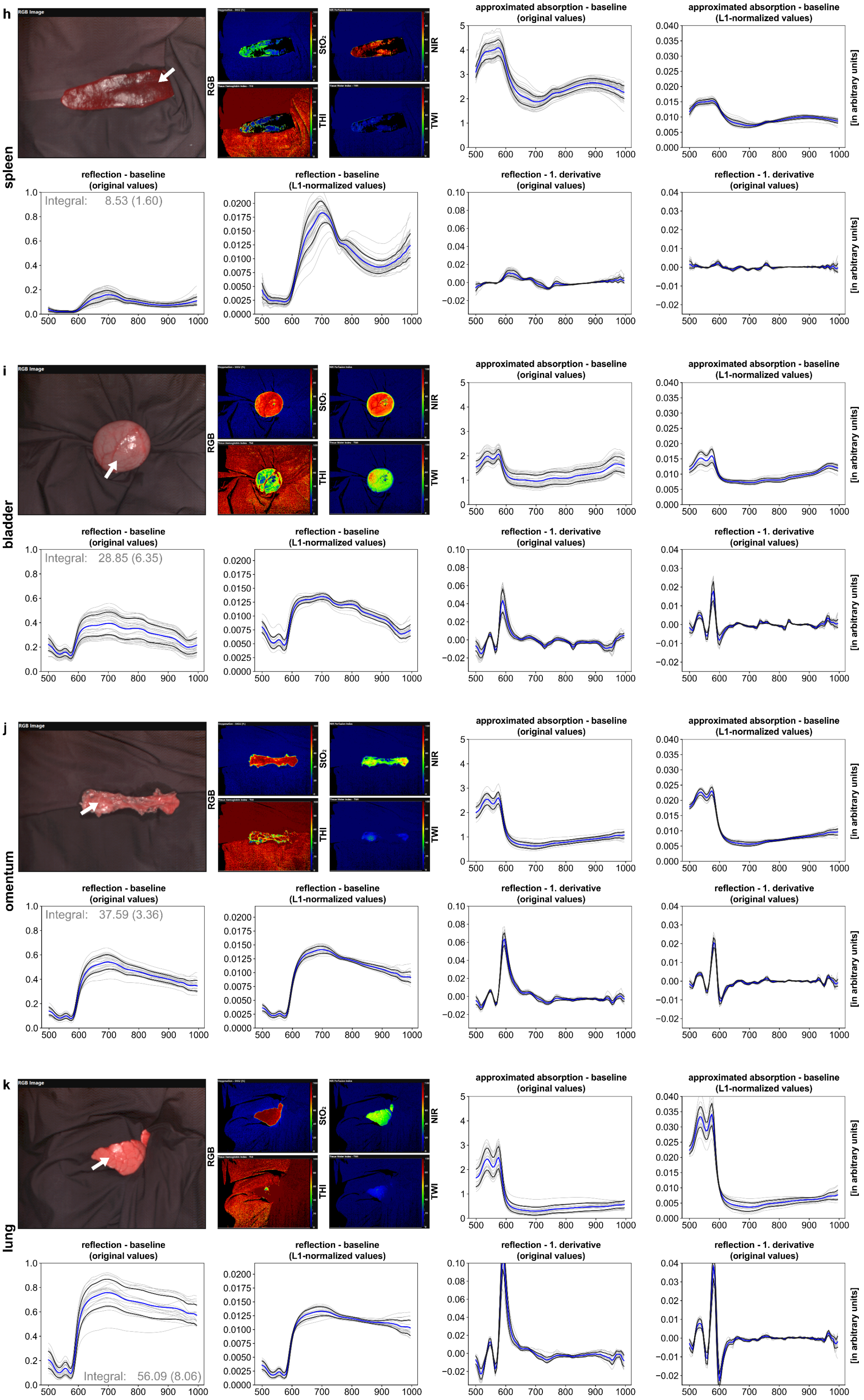
h
spleen
RGB Image
RGB
StO2
NIR
THI
TWI
approximated absorption - baseline (original values)
approximated absorption - baseline (L1-normalized values)
[in arbitrary units]
reflection - baseline (original values)
Integral: 8.53 (1.60)
reflection - baseline (L1-normalized values)
reflection - 1. derivative (original values)
reflection - 1. derivative (original values)
i
bladder
RGB Image
RGB
StO2
NIR
THI
TWI
approximated absorption - baseline (original values)
approximated absorption - baseline (L1-normalized values)
[in arbitrary units]
reflection - baseline (original values)
Integral: 28.85 (6.35)
reflection - baseline (L1-normalized values)
reflection - 1. derivative (original values)
reflection - 1. derivative (original values)
j
omentum
RGB Image
RGB
StO2
NIR
THI
TWI
approximated absorption - baseline (original values)
approximated absorption - baseline (L1-normalized values)
[in arbitrary units]
reflection - baseline (original values)
Integral: 37.59 (3.36)
reflection - baseline (L1-normalized values)
reflection - 1. derivative (original values)
reflection - 1. derivative (original values)
k
lung
RGB Image
RGB
StO2
NIR
THI
TWI
approximated absorption - baseline (original values)
approximated absorption - baseline (L1-normalized values)
[in arbitrary units]
reflection - baseline (original values)
Integral: 56.09 (8.06)
reflection - baseline (L1-normalized values)
reflection - 1. derivative (original values)
reflection - 1. derivative (original values)

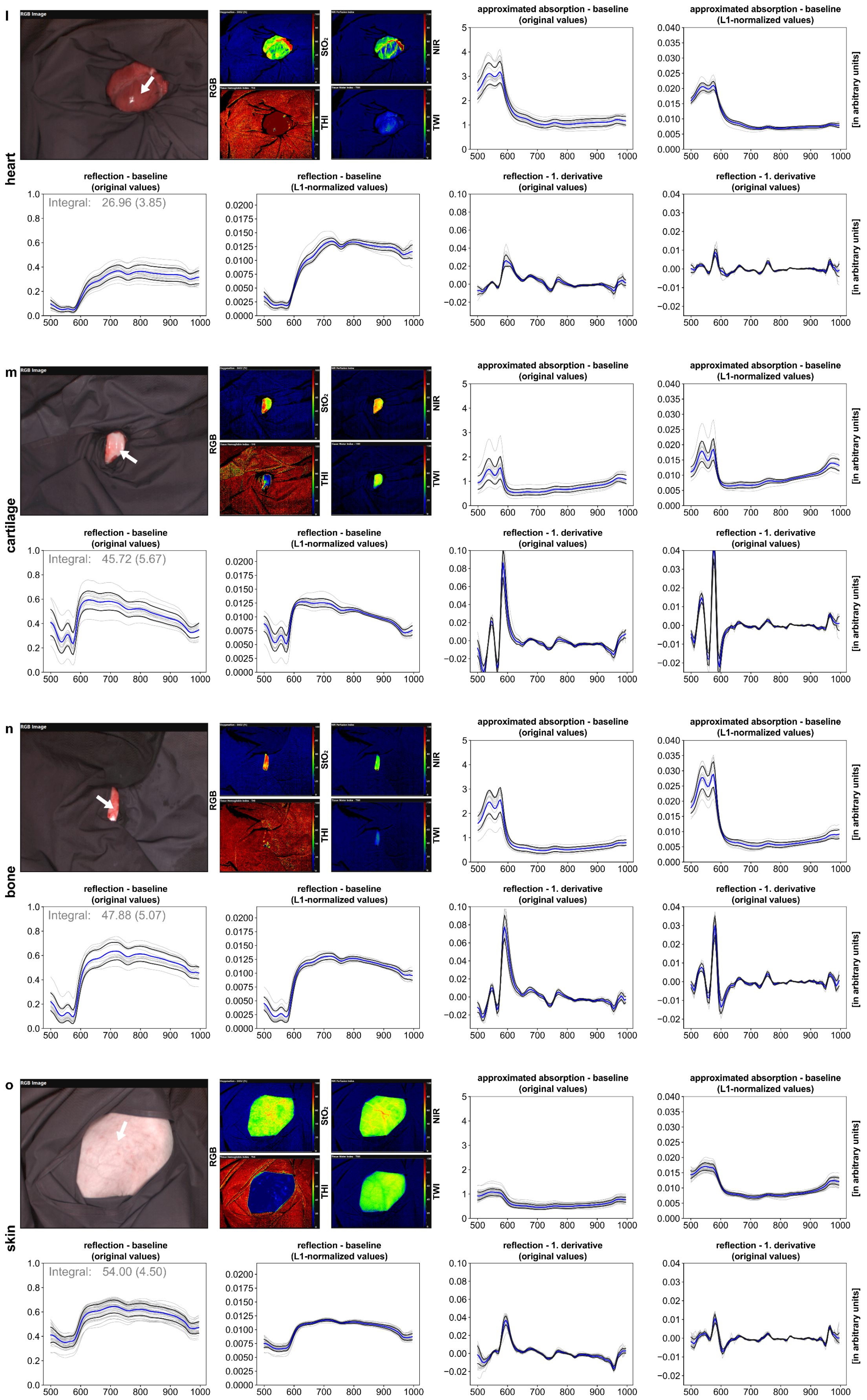
l
heart
RGB Image
RGB
StO₂
NIR
THI
TWI
approximated absorption - baseline (original values)
approximated absorption - baseline (L1-normalized values)
reflection - baseline (original values)
Integral: 26.96 (3.85)
reflection - baseline (L1-normalized values)
reflection - 1. derivative (original values)
[in arbitrary units]
m
cartilage
Integral: 45.72 (5.67)
n
bone
Integral: 47.88 (5.07)
o
skin
Integral: 54.00 (4.50)

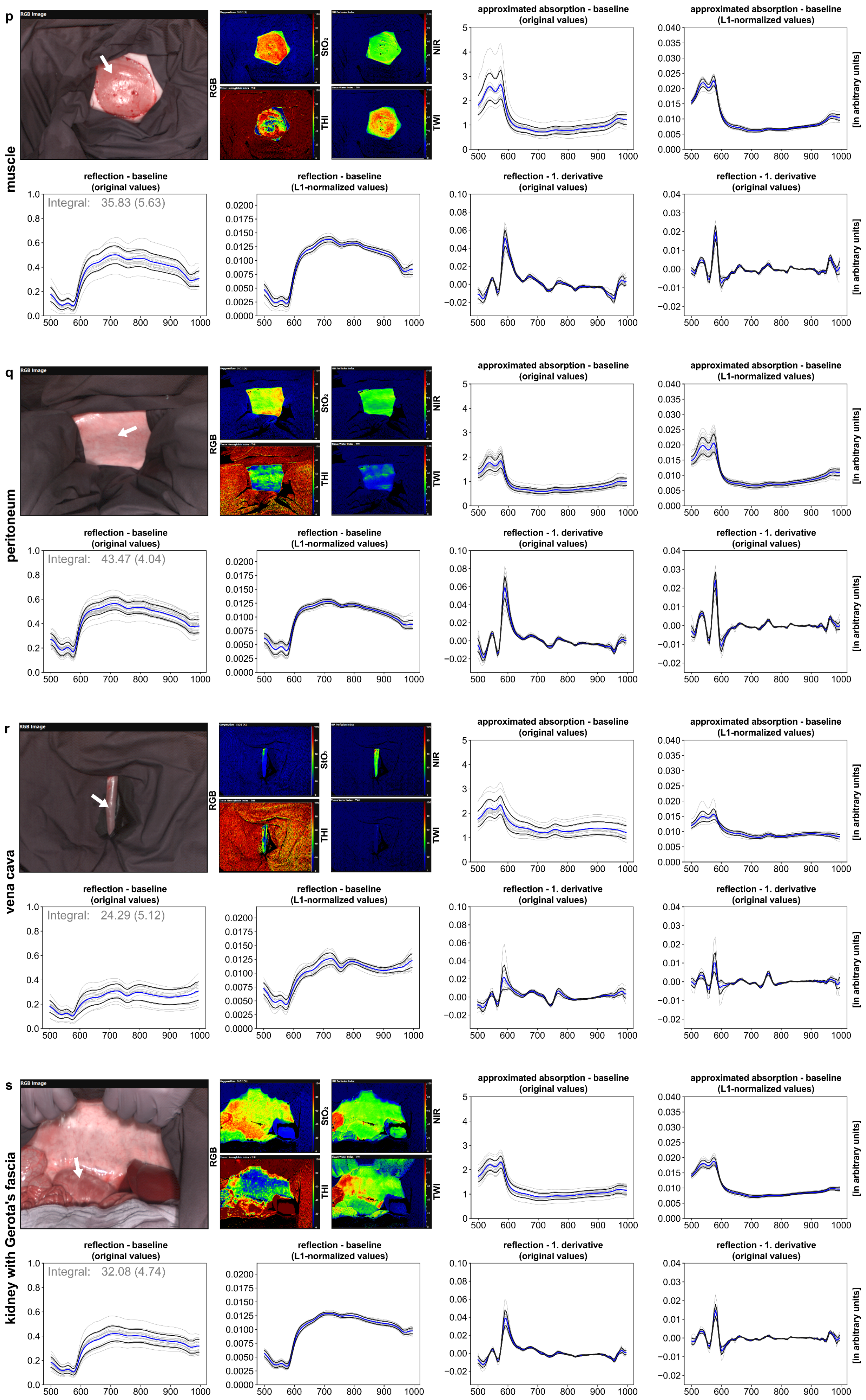
p
muscle
RGB Image
RGB
StO₂
NIR
THI
TWI
approximated absorption - baseline (original values)
approximated absorption - baseline (L1-normalized values)
[in arbitrary units]
reflection - baseline (original values)
Integral: 35.83 (5.63)
reflection - baseline (L1-normalized values)
reflection - 1. derivative (original values)
reflection - 1. derivative (original values)
q
peritoneum
RGB Image
RGB
StO₂
NIR
THI
TWI
approximated absorption - baseline (original values)
approximated absorption - baseline (L1-normalized values)
[in arbitrary units]
reflection - baseline (original values)
Integral: 43.47 (4.04)
reflection - baseline (L1-normalized values)
reflection - 1. derivative (original values)
reflection - 1. derivative (original values)
r
vena cava
RGB Image
RGB
StO₂
NIR
THI
TWI
approximated absorption - baseline (original values)
approximated absorption - baseline (L1-normalized values)
[in arbitrary units]
reflection - baseline (original values)
Integral: 24.29 (5.12)
reflection - baseline (L1-normalized values)
reflection - 1. derivative (original values)
reflection - 1. derivative (original values)
s
kidney with Gerota's fascia
RGB Image
RGB
StO₂
NIR
THI
TWI
approximated absorption - baseline (original values)
approximated absorption - baseline (L1-normalized values)
[in arbitrary units]
reflection - baseline (original values)
Integral: 32.08 (4.74)
reflection - baseline (L1-normalized values)
reflection - 1. derivative (original values)
reflection - 1. derivative (original values)

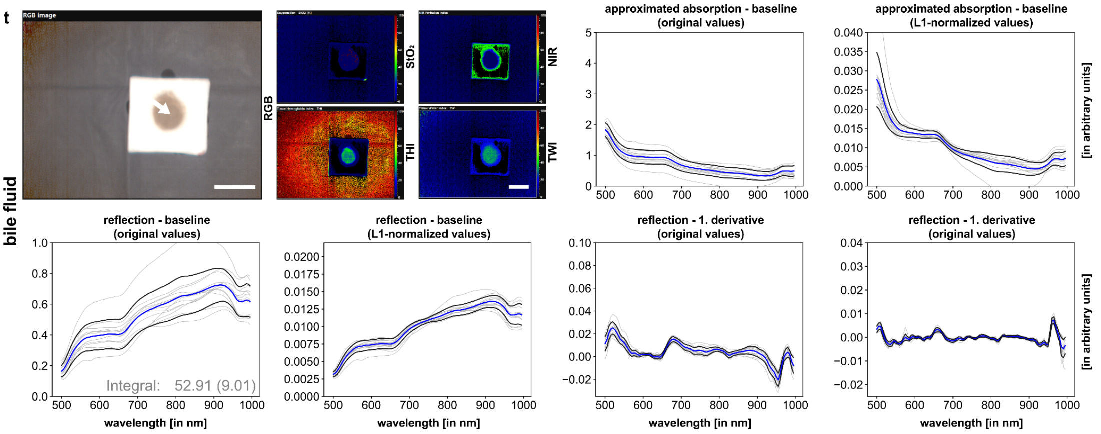


**Supplementary Figure 4 | Hyperspectral tissue characterization for abdominal and thoracic organs.** TIVITA® color-coded images with RGB and index pictures for oxygenation ($StO_2$), perfusion (NIR), tissue hemoglobin (THI) and tissue water (TWI) of representative physiological organs as well as spectral graphs. **a**, stomach. **b**, jejunum. **c**, colon. **d**, liver. **e**, gallbladder. **f**, pancreas. **g**, kidney. **h**, spleen. **i**, bladder. **j**, omentum. **k**, lung. **l**, heart. **m**, cartilage. **n**, bone. **o**, skin. **p**, muscle. **q**, peritoneum. **r**, vena cava. **s**, kidney with Gerota's fascia. **t**, bile fluid. A indicates the number of animals; n indicates the number of measurements in total. Graphs depict approximated absorption and reflectance in original values as well as in L1-normalized values (L1-normalization on pixel-level). Additionally, graphs depict the first and second derivative of reflectance in original values. Individual animals (gray) with overall mean (blue) ±1 SD (black). White arrows indicate areas of measurement.

**Supplementary Text 4: Quantification of organ specificity**

The sum of variances across components other than the factor “organ” at each wavelength (**Figure 4**) averaged across wavelengths can be used as an indicator for the spectral organ-specificity of the respective organ or tissue class. Organs with lower values have more organ-characteristic spectral signatures across observations and vice versa (**Supplementary Table 1**).

***Supplementary Table 1* | Explained variance of reflectance cumulatively for factors "pig", "angle", "image" und "repetition" and averaged across wavelengths**. This is equivalent to the sum of explained variances averaged across wavelengths. A smaller number indicates a more organ-characteristic spectrum. Sorted by increasing values:

| ***organ*** | ***Explained variance cumulatively for factors other than “organ”, averaged across wavelengths*** |
|---:|---|
| *spleen* | 0.00081 |
| *colon* | 0.00189 |
| *heart* | 0.00193 |
| *liver* | 0.00197 |
| *stomach* | 0.00217 |
| *kidney* | 0.00238 |
| *omentum* | 0.00260 |
| *vena cava* | 0.00264 |
| *jejunum* | 0.00304 |
| *bladder* | 0.00322 |
| *kidney with Gerota’s fascia* | 0.00337 |
| *muscle* | 0.00391 |
| *gallbladder* | 0.00456 |
| *peritoneum* | 0.00476 |
| *cartilage* | 0.00538 |
| *bile fluid* | 0.00595 |
| *bone* | 0.00658 |
| *pancreas* | 0.00790 |
| *skin* | 0.00966 |
| *lung* | 0.01289 |

**Supplementary Text 5: Annotation protocol**

Annotations were done on the unprocessed HSI data using the HyperGUI (https://github.com/MIC-Surgery-Heidelberg/HyperGUI_1.0). Possible labels were: “stomach”, “jejunum”, “colon”, “liver”, “gallbladder”, “pancreas”, “kidney”, “spleen”, “bladder”, “omentum”, “lung”, “heart”, “cartilage”, “bone”, “skin”, “muscle”, “peritoneum”, “vena cava”, “kidney with Gerota’s fascia” and “bile fluid”.

Non-semantic annotation was performed with a multi-point selection tool. Areas were selected as ROIs by omitting any areas with artefacts including tissue kinking, shade from the illumination, marginal areas, superficial blood vessels and fat, contamination with dyes or body fluids such as bile fluid, previous manipulation such as contusion or abrasion and possible impairment of perfusion such as thrombosis.

The regions were selected with the aim of including only highly representative areas; therefore, it is guaranteed that analyzed pixels were always 100% representative of the label. Consequently, there are additional adjacent pixels that could have been selected as well, but were not based on the judgement of the annotator and the premise to not include faulty or non-representative pixels under any circumstance. In case of several possible regions that were separated by aforementioned artefact, the largest and most representative area was selected.
